# 2-arm-PEG-oligocations transiently shield the liver sinusoids to mitigate off-target hepatic expression of mRNA lipid nanoparticles

**DOI:** 10.64898/2026.04.29.721537

**Authors:** Anjaneyulu Dirisala, Bhaskar Chatterjee, Le Bui Thao Nguyen, Kazuko Toh, Miki Matsui-Masai, Xueying Liu, Theofilus A. Tockary, Nan Qiao, Junichi Ishikawa, Jumpei Norimatsu, Yuki Mochida, Shigeto Fukushima, Makoto Oba, Kazunori Kataoka, Satoshi Uchida

**Affiliations:** Innovation Center of NanoMedicine (iCONM), Kawasaki Institute of Industrial Promotion, 3-25-14 Tonomachi, Kawasaki-ku, Kawasaki 210-0821, Japan; Department of Advanced Nanomedical Engineering, Medical Research Laboratory, Institute of Integrated Research, Institute of Science Tokyo, 1-5-45 Yushima, Bunkyo-ku, Tokyo 113-8510, Japan; NANO MRNA Co., Ltd., 2-5-1 Atago, Minato-ku, Tokyo 105-6226, Japan; Medical Chemistry, Graduate School of Medical Science, Kyoto Prefectural University of Medicine, 1-5 Shimogamohangi-cho, Sakyo-ku, Kyoto 606-0823, Japan; Pandemic Preparedness, Infection and Advanced Research Center (UTOPIA), The University of Tokyo, 4-6-1, Shirokanedai, Minato-ku, Tokyo 108-0071, Japan

**Keywords:** mRNA delivery, ionizable lipid nanoparticle, hepatic clearance, liver sinusoid, poly(ethylene glycol), oligocation, mRNA vaccine

## Abstract

Ionizable lipid nanoparticles (iLNPs) are powerful platforms for mRNA-based vaccines and immunotherapies; however, their intrinsic liver tropism compromises both safety and efficacy. Off-target hepatic protein expression from delivered mRNA raises safety concerns, and hepatic clearance limits efficient iLNP delivery to target organs. In this study, we address these challenges in mouse models by stealth-coating the liver sinusoidal endothelial (LSE) wall, the primary gateway for nanoparticle entry into the liver. Specifically, oligocations conjugated with two-armed PEG (2-arm-PEG-oligocations), a clinically relevant material used in oligonucleotide delivery trials, were employed to transiently anchor PEG to the LSE wall with balanced affinity, ensuring robust coating followed by gradual biliary clearance. This approach reduced hepatic protein expression from iLNPs, subsequently administered either systemically or locally, by more than tenfold. Importantly, the strategy preserved iLNP accumulation in the spleen, a key target organ for vaccines, effectively redirecting iLNPs from the liver to the spleen. Consequently, in vaccine applications, pre-injection of the 2-arm-PEG-oligocation preserved or even enhanced vaccination efficacy while minimizing concerns associated with antigen expression in the liver. In applications involving cytokine mRNA therapy, specifically intratumoral interleukin-12 (IL-12) mRNA administration, systemic pre-injection of the 2-arm-PEG-oligocation successfully reduced off-target hepatic IL-12 expression and subsequent systemic IL-12 exposure, while maintaining antitumor efficacy. Collectively, these results demonstrate that LSE-wall stealth coating is a generalizable strategy to improve both the safety and efficacy of iLNP-based mRNA vaccines and immunotherapies.

## INTRODUCTION

mRNA vaccines demonstrated clear clinical utility during the COVID-19 pandemic,^1, 2^ catalyzing extensive research and development of mRNA-based applications across diverse medical fields, including cancer immunotherapy, protein replacement therapy, and genome editing.^3, 4^ Among these applications, vaccines and immunotherapies have been studied intensively and are now advancing into clinical development. Nanoparticle-based mRNA delivery technologies, together with mRNA engineering approaches such as nucleoside modification,^5–7^ have substantially contributed to this progress, with ionizable lipid nanoparticles (iLNPs) recently garnering particular attention.^8–10^ In addition to protecting mRNA from nucleases and facilitating endosomal escape, iLNPs possess additional functionalities that are advantageous for vaccines and immunotherapies.^11, 12^ Their intrinsic immunostimulatory properties act as built-in adjuvants in vaccines^13, 14^ and potentially enhance the efficacy of cancer immunotherapy by activating pro-inflammatory responses. Effective targeting of lymphoid tissues is another essential feature of iLNPs, particularly for vaccine applications. iLNPs efficiently accumulate in draining lymph nodes after local injection,^11, 12, 15, 16^ and can be engineered to target the spleen, the largest secondary lymphoid organ, following systemic administration.^17–22^ When antigen presenting cells (APCs) in these lymphoid organs express antigens encoded by mRNA, they effectively promote immunization through interactions with neighboring T and B cells.^12, 23–25^

Despite these advantages, iLNPs also accumulate in off-target organs, most notably the liver, through multiple mechanisms. Apolipoprotein E (ApoE) binds to iLNPs in the bloodstream and promotes their uptake by hepatocytes via low-density lipoprotein receptors (LDLRs).^26–28^ In addition, liver sinusoidal endothelial cells (LSECs) and Kupffer cells (KCs) exhibit high scavenging activity and efficiently capture nanoparticles.^29–31^ The slow blood flow within liver sinusoids further enhances this uptake.^32^ Notably, iLNPs accumulate in the liver even after local administration, such as intramuscular or intratumoral injection, due to leakage into systemic circulation from the injection site.^33–39^

Hepatic accumulation of iLNPs can compromise both the efficacy and safety of vaccines and immunotherapies via two major mechanisms. First, hepatic clearance of iLNPs reduces their availability in target organs, such as the spleen in vaccine applications, thereby diminishing therapeutic efficacy. Second, unintended protein expression in the liver raises safety and efficacy concerns. Pro-inflammatory responses to iLNPs in the liver are amplified, particularly under pre-existing inflammatory conditions.^40^ In cancer immunotherapy involving intratumoral delivery of cytokine mRNA,^41–45^ ectopic cytokine expression in the liver may trigger pro-inflammatory responses that can propagate systemically.^34, 39^ In vaccines, unintended antigen expression in the liver can induce autoimmune responses, as suggested by rare cases of autoimmune hepatitis reported in humans following COVID-19 mRNA vaccination.^46, 47^ A mechanistic study revealed enrichment of cytotoxic T lymphocytes (CTLs) targeting the spike protein in the liver of an affected patient, suggesting that these CTLs may attack liver cells expressing the antigen.^48^ Conversely, hepatic accumulation of iLNPs could also induce immune tolerance and counteract vaccine efficacy due to the tolerogenic properties of non-parenchymal liver cells, including KCs and LSECs.^49, 50^ Indeed, a previous study successfully modulated peanut allergy by delivering mRNA iLNPs encoding peptide epitopes to LSECs.^51^

Blocking the liver sinusoidal endothelial (LSE) wall, the primary entry point for nanoparticles into the liver, represents a potential strategy to reduce hepatic accumulation of iLNPs. Because liver uptake is a common challenge in nanomedicine, several approaches have been explored to block the LSE wall, including pre-injection of scavenger receptor ligands^52–54^ or decoy nanoparticles.^55–57^ However, these strategies suffer from limitations in both safety and efficacy. Many scavenger receptor ligands raise toxicological concerns,^58–60^ and a prior study has shown that pretreatment with scavenger receptors-blocking agents, such as dextran sulfate or decoy liposomes, fails to prevent hepatic clearance of iLNPs, underscoring the difficulty of this approach.^61^ While most LSE wall-blocking strategies are designed to inhibit specific nanoparticle uptake pathways, iLNPs accumulate in the liver through multiple mechanisms, including LDLR-mediated uptake of ApoE-bound particles,^26–28^ sequestration by LSECs and KCs,^29–31^ and slow sinusoidal blood flow.^32^ Consequently, effective inhibition of hepatic iLNP clearance likely requires strategies capable of broadly suppressing these pathways simultaneously.

In this context, stealth coating of the LSE wall using poly(ethylene glycol) (PEG) offers a promising approach to broadly inhibit iLNP binding without relying on specific uptake mechanisms. Indeed, a previous study has demonstrated that this strategy effectively suppresses liver accumulation of diverse nanoparticle platforms, including viral vectors and polymeric nanoparticles, suggesting its broad potential to inhibit multiple mechanisms of hepatic clearance.^62^ In this approach, two-armed PEG is conjugated to an oligocation, enabling PEG anchoring to the LSE wall through interactions with anionic proteoglycans or scavenger receptors. Systemic injection of the resulting 2-arm-PEG-oligocation enables transient and selective PEG coating of the LSE wall,^62, 63^ which is essential for avoiding prolonged disruption of liver function and minimizing effects on other organs. In contrast, the 1-arm-PEG-oligocation formulation persists within the LSE wall for extended periods, raising potential safety concerns. Mechanistic analyses revealed that 2-arm-PEG-oligocation, but not 1-arm-PEG-oligocation, is efficiently excreted into bile within several hours^62^. Notably, 2-arm-PEG-oligocation is currently being evaluated in phase I clinical trials as a delivery carrier for small interfering RNA (siRNA) or antisense oligonucleotides (ASOs),^64–66^ and has demonstrated negligible toxicity in rigorous preclinical studies,^67^ underscoring its clinical translatability.

In this study, we demonstrate in mice that PEG coating of the LSE wall dramatically reduces liver accumulation of iLNPs and subsequent hepatic protein expression from delivered mRNA (**Scheme 1**). Importantly, this strategy does not inhibit iLNP accumulation in the spleen, a critical organ for mRNA vaccination, and instead enhances splenic protein expression efficiency after systemic administration by redirecting iLNPs from the liver to the spleen. As a result, the vaccination efficacy of systemically or locally administered iLNPs is preserved or even enhanced following LSE wall coating, while off-target protein expression in the liver is markedly reduced. Furthermore, LSE coating also improves the safety of cancer immunotherapies involving intratumoral administration of cytokine mRNA. Although intratumorally injected iLNPs leak into systemic circulation and induce cytokine expression in the liver, PEG coating reduces hepatic cytokine expression without compromising antitumor efficacy. Together, these findings highlight the potential of LSE-wall PEG coating as a broadly applicable strategy to improve both the safety and efficacy of mRNA vaccines and immunotherapies.

**Scheme 1.**
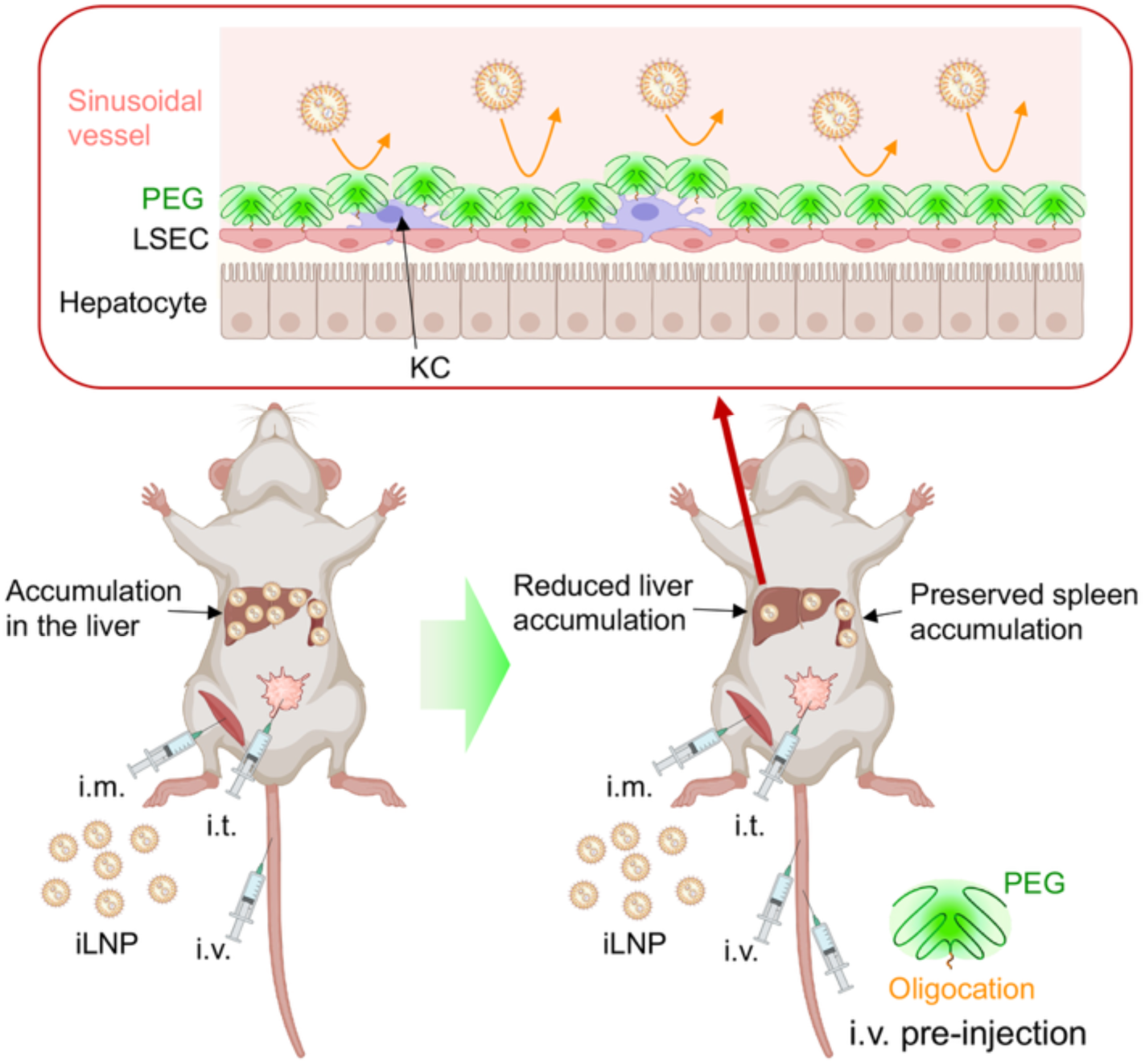
Concept of this study. Ionizable lipid nanoparticles (iLNPs) typically accumulate in the liver following systemic intravenous (i.v.) administration as well as local administrations such as intramuscular (i.m.) and intratumoral (i.t.) injection. This study introduces a strategy to inhibit hepatic accumulation of iLNPs by coating the liver sinusoidal endothelial (LSE) wall with poly(ethylene glycol) (PEG). For PEG coating, a 2-arm PEG is conjugated to an oligocation, enabling anchoring to the LSE wall. Pre-injection of the 2-arm PEG–oligocation effectively reduces liver accumulation of iLNPs while preserving their accumulation in the spleen, an important target organ for mRNA vaccines. LSEC: liver sinusoidal endothelial cell. KC: Kupffer cell. Prepared using BioRender under license.

## RESULTS AND DISCUSSIONS

### 2-arm-PEG-oligocations reduce iLNP accumulation in the liver but not in the spleen

We selected ALC-0315-based iLNPs, which are used in the approved COVID-19 mRNA vaccine, BNT162b2, due to their strong potential for mRNA vaccination and delivery.^1, 2, 38^ The iLNPs exhibited a hydrodynamic diameter of 83.8 ± 0.7 nm and a polydispersity index (PDI) of 0.09 ± 0.02 as determined by dynamic light scattering (DLS), a ζ-potential of – 5.8 ± 1.7 mV measured by laser Doppler electrophoretic light scattering, and an encapsulation efficiency of 95.7 ± 0.8 % (mean ± standard deviation; **Table S1**).

For PEG coating of the LSE wall, two-armed PEG with 40 kDa PEG chains in each arm was conjugated to an oligocationic peptide consisting of approximately 20 repeating units. A previous study demonstrated that the large PEG chains enhance the selectivity of oligocation accumulation in the LSE wall by preventing attachment to extrahepatic endothelia, presumably due to steric repulsion by PEG.^62^ At the same time, the oligocation effectively accumulates in the LSE wall even after 2-arm PEG conjugation, likely owing to the high abundance of heparan sulfate proteoglycans and scavenger receptors in this endothelium, which strongly attract oligocations. In the present study, oligolysine (OligoLys) or oligoornithine (OligoOrn) was used as the oligocation (**Scheme 2**), both of which have demonstrated high safety and efficacy in oligonucleotide delivery when formulated as 2-arm-PEG-oligocations.^63, 66, 67^ Notably, these two oligocation species differ only in the length of the side-chain spacer: OligoLys possesses a tetramethylene spacer, whereas OligoOrn contains a trimethylene spacer. Among these candidates, we initially selected OligoLys for the cationic segment, as a previous study thoroughly elucidated the intrahepatic behavior of 2-arm-PEG-OligoLys.^62^

**Scheme 2.**
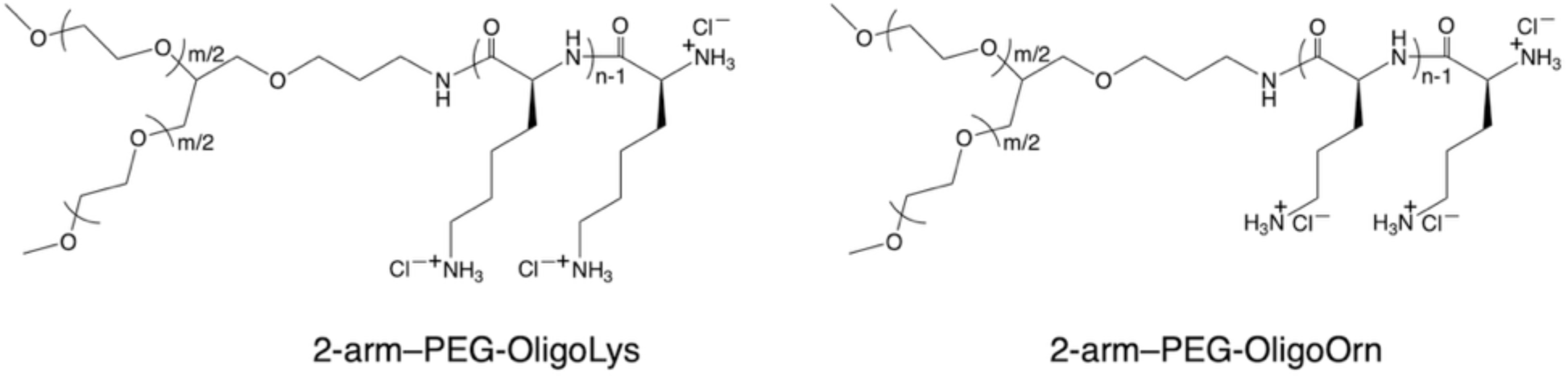
Chemical structures of 2-arm-PEG-oligocations.

In developing a strategy that combines LSE wall stealth coating with iLNP delivery, we considered the possibility that the 2-arm-PEG-oligocation might interact with iLNPs in the bloodstream, thereby affecting their mRNA delivery performance. To investigate this possibility, we evaluated the diffusion properties of 2-arm-PEG-OligoLys in solution using fluorescence correlation spectroscopy (FCS). **Table 1** shows the diffusion time of Alexa647-labeled 2-arm-PEG-OligoLys, which is inversely correlated with the diffusion coefficient and would increase if the 2-arm-PEG-oligocation interacted with ALC-0315-based iLNPs. Notably, the diffusion time remained unchanged in the presence of iLNPs, indicating minimal interaction between the 2-arm-PEG-oligocation and the iLNPs.

**Table 1.** Diffusion properties of 2-arm-PEG-OligoLys.

|  | iLNP (-) | iLNP (+) |
| --- | --- | --- |
| Diffusion time (μsec) | $270.2 \pm 17.7$ | $286.5 \pm 32.3$ |

Fluorescence correlation spectroscopy (FCS) analysis of Alexa647-labeled 2-arm-PEG-OligoLys in the presence or absence of iLNPs. Data are presented as mean ± standard error of the mean. $n = 11$ for iLNP (-), $n = 18$ for iLNP (+)

In animal studies, we first examined the biodistribution of the 2-arm-PEG-oligocation using Alexa594-labeled 2-arm-PEG-OligoLys. Thirty minutes after systemic injection in mice, the 2-arm-PEG-oligocation predominantly accumulated in the liver (**Figure 1A**). Regarding intrahepatic distribution, previous work demonstrated that the 2-arm-PEG-oligocation attaches to the LSE wall within 5 min and remains there for several hours following systemic administration.^62^ With respect to extrahepatic distribution, smaller amounts of the 2-arm- PEG-oligocation were detected in the spleen and kidney, whereas fluorescence signals were nearly undetectable in the lung, heart, and brain (**Figure 1A**).

**Figure 1.**
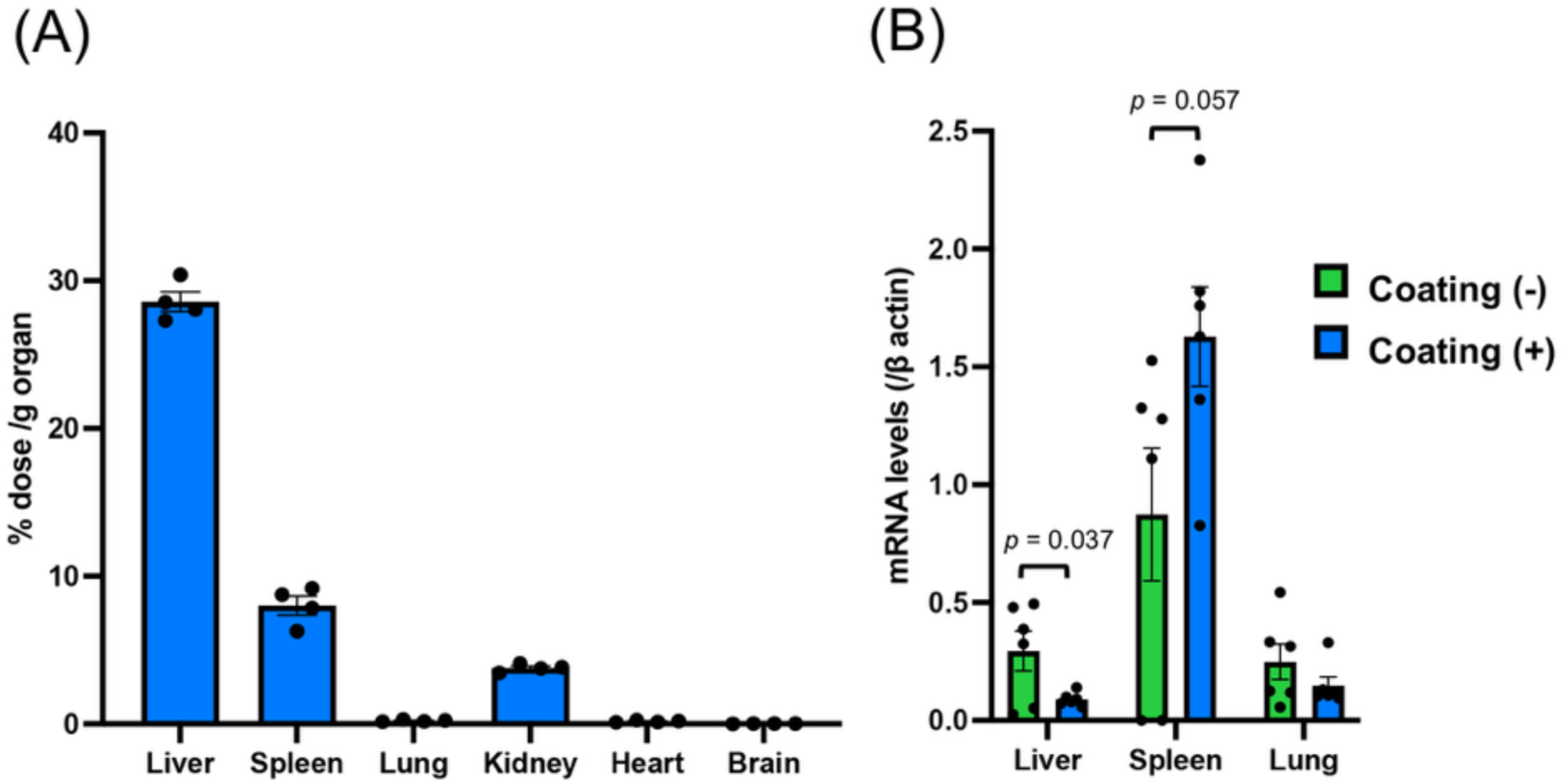
Biodistribution of PEGylated oligocations and iLNPs in mice. (A) Alexa594-labeled 2-arm-PEG-OligoLys was intravenously injected, and organ distribution was quantified 30 min post-injection by measuring fluorescence intensity in tissue homogenates. *n* = 4. (B) ALC-0315-based iLNPs were intravenously injected with or without pre-injection of 2-arm-PEG-OligoLys 5 min beforehand. iLNP biodistribution was quantified 4 h post-injection by qPCR analysis of delivered mRNA. *n* = 6. Data are presented as mean ± SEM. Statistical analyses were performed using an unpaired two-tailed Student’s *t*-test.

Next, we evaluated the ability of the 2-arm-PEG-oligocation to modulate iLNP biodistribution. For LSE wall stealth coating, the 2-arm-PEG-oligocation was systemically injected five minutes prior to systemic administration of the iLNPs. Four hours after iLNP injection, biodistribution was assessed by quantitative PCR (qPCR) analysis of delivered mRNA in each organ. Pre-injection with the 2-arm-PEG-oligocation significantly reduced mRNA levels in the liver compared with the control group without pre-injection (**Figure 1B**), demonstrating that the 2-arm-PEG-oligocation effectively prevents iLNP accumulation in the liver. In contrast, in the spleen, pre-injection with the 2-arm-PEG-oligocation did not reduce iLNP accumulation, despite the noticeable accumulation of the 2-arm-PEG-oligocation in this organ (**Figure 1A**). In the lung, pre-injection had little impact on iLNP accumulation, consistent with the minimal oligocation accumulation observed in this organ. In other examined organs, including the kidney, heart, and brain, mRNA levels were close to background, precluding reliable quantification.

### 2-arm-PEG-oligocations reduce LSE wall attachment and parenchymal migration of iLNPs in the liver

To investigate the behavior of iLNPs at the microscopic level, we visualized 2-arm-PEG-oligocations and iLNPs using intravital real-time confocal laser-scanning microscopy (IVRT-CLSM) following intravenous injection. First, the liver was imaged after administration of DiD-labeled ALC-0315-based iLNPs (red), with or without pre-injection of Alexa555-labeled 2-arm-PEG-OligoLys (cyan) 5 min beforehand (**Figure 2A**). Hepatocyte autofluorescence was used to visualize the liver parenchyma (green), thereby delineating sinusoidal vessels as regions lacking green fluorescence. Thirty minutes after iLNP injection in the absence of 2- arm-PEG-oligocation pre-injection, strong red fluorescent signals were observed on both sides of the vessel structures, indicating substantial accumulation of iLNPs along the LSE wall (**Figure 2B**). These signals are indicated by white arrowheads in **Figure 2C** at the 30-min time point. One hour or later after injection, red fluorescent iLNPs progressively migrated deeper into the liver parenchyma, resulting in expanded areas of yellow pixels, as highlighted by the dotted circles in **Figure 2C**, indicating colocalization of iLNPs with hepatocyte autofluorescence.

**Figure 2.**
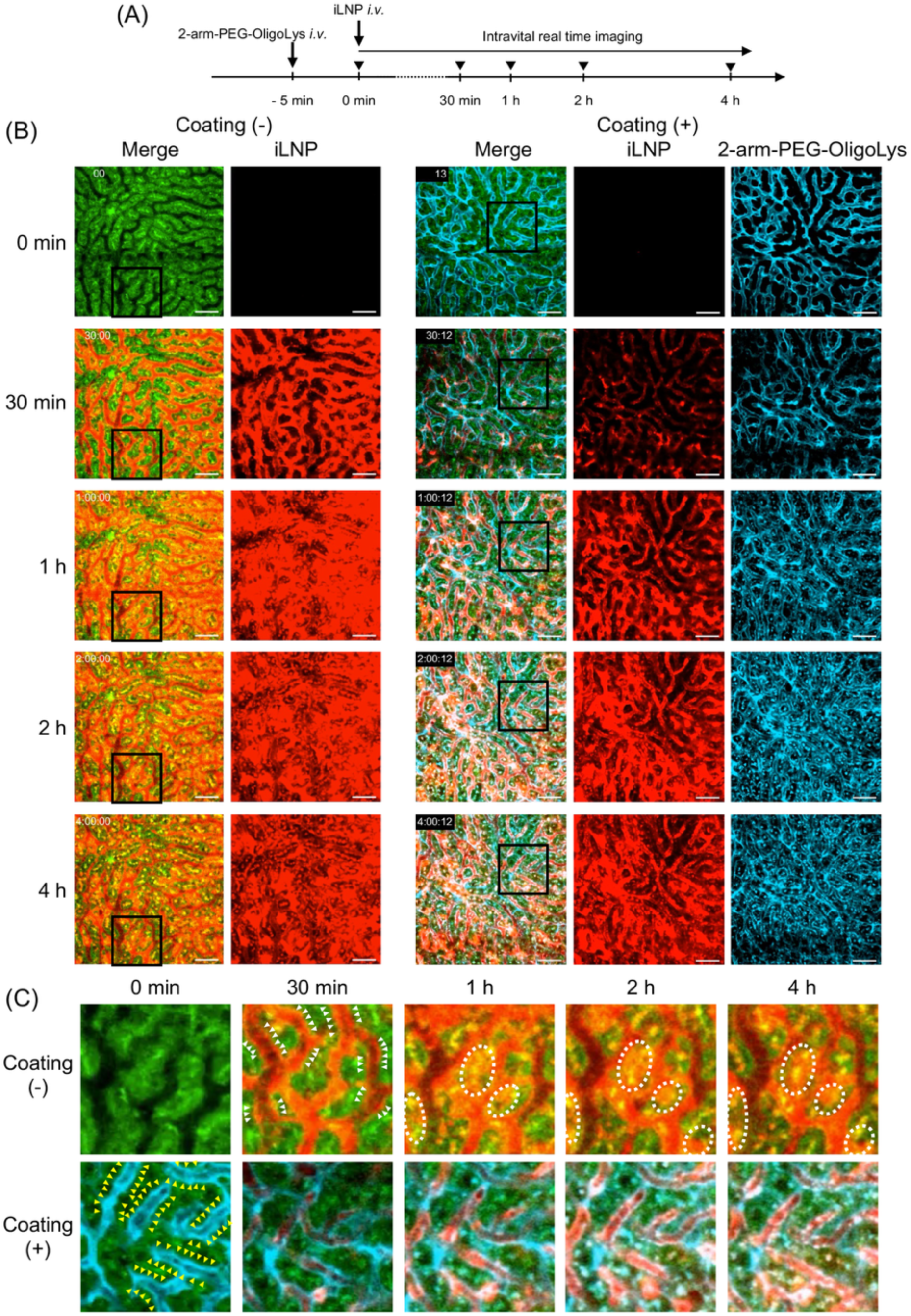
Real-time observation of 2-arm-PEG-oligocations and iLNPs in the liver. Alexa555-labeled 2-arm-PEG-OligoLys (cyan) and DiD-labeled ALC-0315-based iLNPs (red) were intravenously injected into mice at a 5 min interval (Coating (+)). The control group (Coating (-)) did not receive 2-arm-PEG-OligoLys. Liver parenchymal autofluorescence is shown in green. (A) Experimental timeline. (B) Images acquired at the indicated time points after iLNP injection. (C) Enlarged views of the boxed regions in (B). White arrowheads indicate iLNP attachment to the LSE wall. Dotted circles indicate migration of iLNPs into the liver parenchyma. Yellow arrowheads indicate attachment of 2-arm-PEG-OligoLys to the LSE wall.

In the 2-arm-PEG-oligocation pre-injection group, strong cyan fluorescence signals were observed along the lining of both sides of the vessel structures at 5 min post-injection (defined as 0 min in **Figure 2B, C**), as indicated by yellow arrowheads in **Figure 2C**. This observation indicates successful PEG coating of the LSE wall. In contrast to the group without pre-injection, accumulation of red-fluorescent iLNPs along the sinusoidal walls was not apparent at 30-min after iLNP administration in the presence of the 2-arm-PEG-oligocation. This finding indicates that PEG coating of the LSE wall effectively inhibits iLNP attachment. Consequently, pre-injection of the 2-arm-PEG-oligocation markedly suppressed subsequent migration of iLNPs into the liver parenchyma at later time points, as evidenced by reduced colocalization of red (iLNP) and green (parenchymal) fluorescence in **Figure 2B, C**. These microscopic observations are consistent with the qPCR-based biodistribution analysis (**Figure 1B**), collectively indicating that LSE wall stealth coating effectively reduces hepatic accumulation of iLNPs.

### iLNPs migrate into the splenic parenchyma despite accumulation of 2-arm-PEG-oligocations

In contrast to the liver, pre-injection of the 2-arm-PEG-oligocation did not inhibit iLNP migration into the spleen (**Figure 1B**), despite substantial accumulation of the 2-arm-PEG-oligocation in this organ (**Figure 1A**). To investigate the underlying mechanisms, we performed intravital imaging of Alexa450-labeled 2-arm-PEG-OligoLys (green) and DiD-labeled ALC-0315-based iLNPs (red) in the spleen (**Figure 3A**). The 2-arm-PEG-oligocation was distributed throughout the field of view in the spleen shortly after injection (**Figure 3B**). Notably, luminal structures composed of strong green-fluorescence were observed, with signal enrichment along both sides of these structures, as outlined by dotted lines in the enlarged images (rightmost panels in **Figure 3B**). These observations suggest that PEG may also coat splenic endothelial walls, in addition to the LSE wall. Despite the widespread presence of the 2-arm-PEG-oligocation in the spleen, red fluorescent iLNPs became broadly distributed throughout the splenic tissue within 30 min. Time-resolved imaging revealed a heterogeneous distribution of iLNPs at early time points: five minutes after injection, iLNPs were confined to localized regions of the spleen, but their distribution progressively expanded over time. This pattern suggests that iLNPs might enter the spleen through specific anatomical sites prior to dispersal throughout the parenchyma.

**Figure 3.**
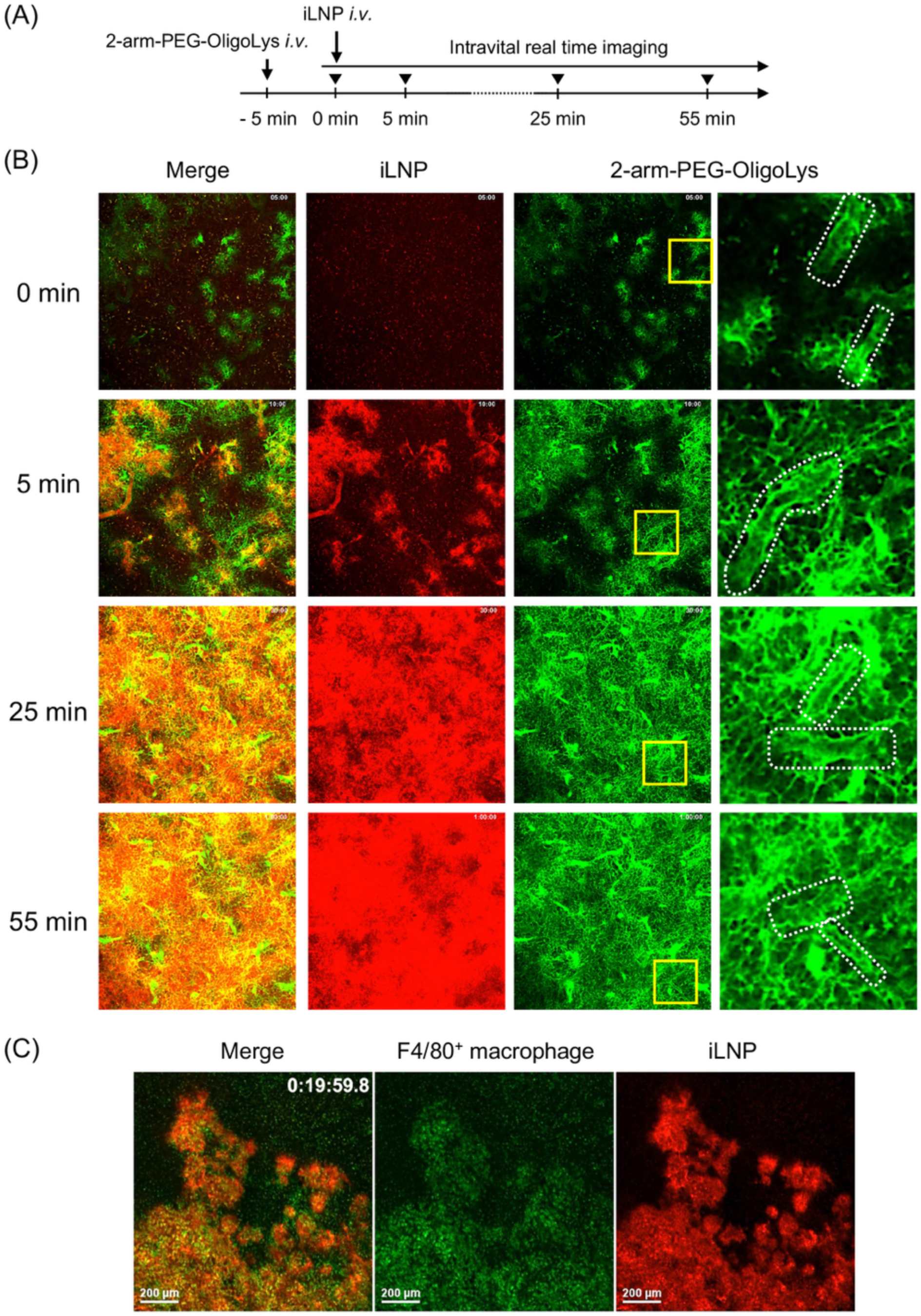
Real-time observation of PEGylated oligocations and iLNPs in the spleen. (A, B) Alexa450-labeled 2-arm-PEG-OligoLys (green) and DiD-labeled ALC-0315-based iLNPs (red) were intravenously injected at a 5 min interval. (A) Experimental timeline. (B) Images acquired at the indicated time points after iLNP injection. The rightmost images show magnified views of the areas indicated by the yellow squares in the second images from the right. Luminal structures are outlined by dotted lines. (C) F4/80-positive macrophages were labeled by intravenous injection of Alexa488-conjugated anti-F4/80 antibody (green). Mice were subsequently injected with unlabeled 2-arm-PEG-OligoLys followed by DiD-labeled iLNPs (red) at a 5-min interval. Images were acquired 10 min after iLNP injection.

Notably, the spleen contains both closed and open circulatory pathways, with the open circulation allowing blood to flow directly into the red pulp parenchyma without passage through continuous endothelial linings.^68, 69^ This open circulatory system might enable iLNPs to access the splenic parenchyma even when splenic blood vessels are coated with PEG. To support this hypothesis, we visualized the red pulp by labeling F4/80 positive macrophages, predominantly localized within this region,^70^ using a pre-injected Alexa488-labeled anti-F4/80 antibody (**Figure 3C**). Subsequently, unlabeled 2-arm-PEG-oligocation and DiD-labeled iLNPs were injected sequentially with a five-minute interval. Ten minutes after iLNP injection, iLNPs were primarily detected in areas enriched with F4/80 positive cells, indicating that iLNPs predominantly enter the spleen via the red pulp. This pathway could account for the efficient accumulation of iLNPs in the spleen, despite the abundance of the 2-arm-PEG-oligocation in this organ.

### Stealth coating of the LSE wall shifts protein expression distribution from the liver to the spleen

Next, we evaluated the organ distribution of protein expression following administration of mRNA-loaded iLNPs using *luciferase* mRNA as a reporter, measured 4 h after iLNP injection. Regardless of 2-arm-PEG-oligocation pre-injection, the liver and spleen were the two primary organs exhibiting luciferase expression (**Figure 4A**). Notably, 2-arm-PEG-oligocation pre-injection reduced luciferase expression in the liver by 22-fold following systemic iLNP administration, while significantly increasing expression efficiency in the spleen by 2.3-fold. When normalized to organ weight, the ratio of luciferase expression in the spleen relative to the liver increased from 0.13 without pre-injection to 6.1 with pre-injection (**Figure 4B**). Overall, this corresponds to an approximately 50-fold enhancement in spleen selectivity.

**Figure 4.**
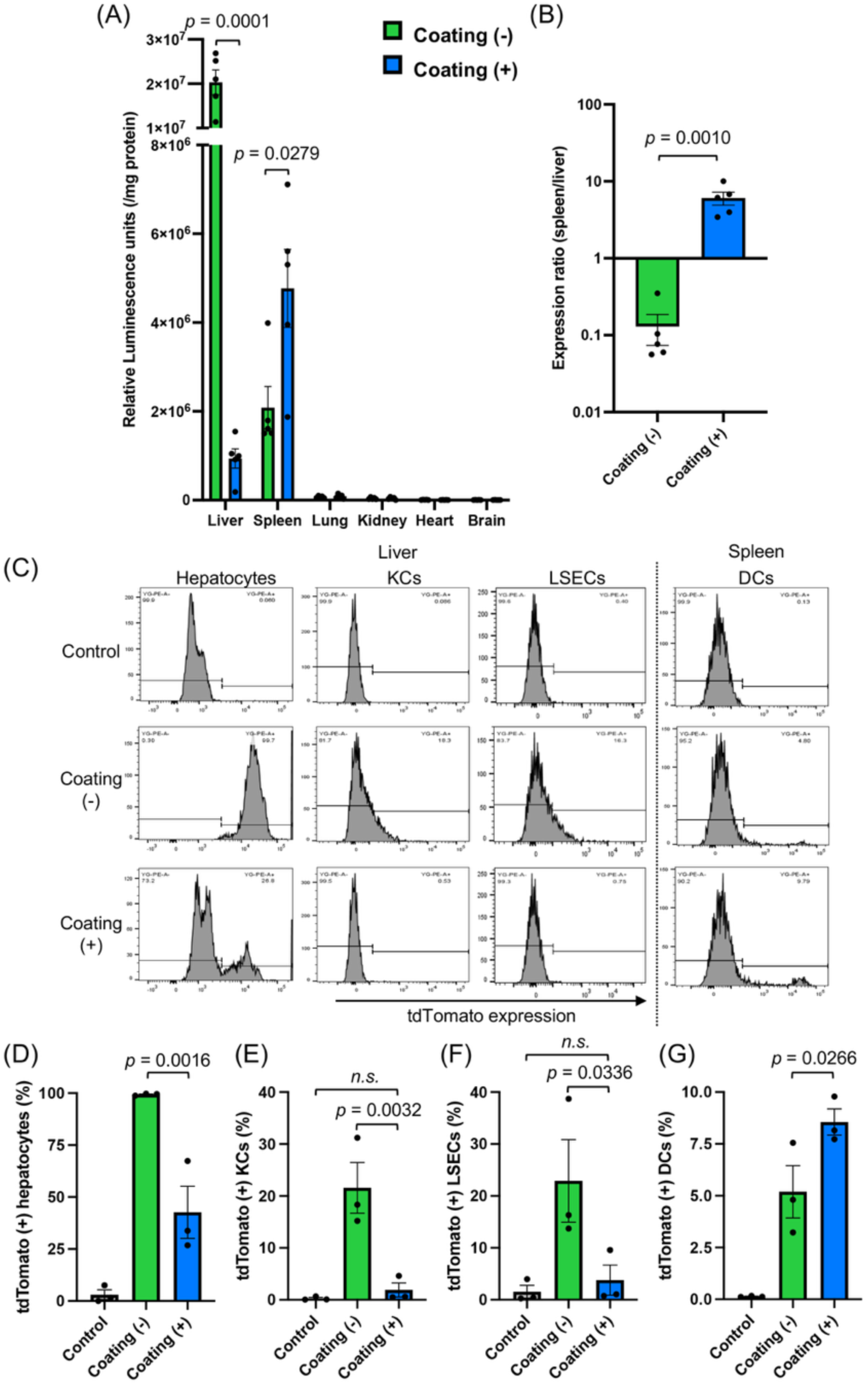
Effects of LSE wall stealth coating on protein expression from delivered mRNA. (A, B) ALC-0315-based iLNPs encapsulating *luciferase* mRNA were intravenously injected with or without pre-injection of 2-arm-PEG-OligoLys 5 min beforehand. Luciferase expression in organ homogenates was quantified 4 h post-injection of iLNPs. (A) Luciferase expression levels in individual organs. (B) Relative luciferase expression in the spleen compared with the liver, normalized to organ weight. *n* = 5. (C-G) ALC-0315-based iLNPs encapsulating *Cre* mRNA were intravenously injected into Ai9 mice with or without 2-arm-PEG-OligoLys pre-injection 5 min beforehand. tdTomato expression was analyzed by flow cytometry 3 days later in hepatocytes (D), Kupffer cells (KCs) (E), liver sinusoidal endothelial cells (LSECs) (F), and splenic dendritic cells (DCs) (G). *n* = 3. Data are mean ± SEM. Statistical analyses used an unpaired two-tailed Student’s t test (A, B) and one-way ANOVA followed by Fisher’s least significant difference post hoc test (D-G).

This protein expression pattern was consistent with iLNP biodistribution assessed by qPCR analysis of delivered mRNA (**Figure 1B**). PEG coating of the LSE wall significantly reduced mRNA levels in the liver and showed a trend toward increased mRNA levels in the spleen, although the latter did not reach statistical significance (p = 0.057). These observations likely reflect the redirection of iLNPs from the liver to the spleen after PEG coating. In the lungs, which exhibited the third-highest level of luciferase expression after the liver and spleen, 2-arm-PEG-oligocation pre-injection had no measurable effect on luciferase expression levels (**Figures 4A, S1**). This finding is consistent with minimal accumulation of the 2-arm-PEG-oligocation in the lung (**Figure 1A**) and the negligible effect of pre-injection on iLNP accumulation in this organ (**Figure 1B**). Pre-injection also did not significantly affect expression levels in the, kidney, heart, or brain, presumably due to limited PEG coating of the endothelium in these organs as can be seen from **Figure 1A**.

A similar trend was observed at 24 h post-injection, although overall expression levels were lower than those measured at 4 h (**Figure S2**). The spleen-to-liver expression ratio increased from 0.78 without 2-arm-PEG-oligocation pre-injection to 21 with pre-injection, corresponding to a 28-fold enhancement in spleen selectivity. Safety analyses revealed no elevations in markers of liver injury (AST, ALT), kidney function (BUN, creatinine), or systemic tissue damage (LDH) 24 h after iLNP injection, with or without 2-arm-PEG-oligocation pre-injection (**Table S2**).

We next examined protein expression at the cellular level in the liver and spleen. *Cre* mRNA was administered to Ai9 reporter mice, which express the fluorescent protein tdTomato upon Cre-mediated recombination at *loxP* sites.^71^ Flow cytometric analysis performed 3 days post-injection revealed that, in the absence of 2-arm-PEG-oligocation pre-injection, iLNPs induced tdTomato expression in nearly 100% of hepatocytes and in approximately 20% of F4/80-positive KCs and CD146-positive LSECs (**Figure 4C-F**). Pre-injection with the 2-arm-PEG-oligocation significantly reduced the percentages of tdTomato-positive hepatocytes, KCs, and LSECs, with expression in KCs and LSECs reduced to near-background levels. In contrast, analysis of CD11c-positive dendritic cells (DCs) in the spleen, key cellular targets in systemic mRNA vaccination, revealed that 2-arm-PEG-oligocation pre-injection increased the percentage of tdTomato positive DCs (**Figure 4C, G**). This result is consistent with the enhanced luciferase expression observed in the spleen following LSE wall stealth coating (**Figure 4A**).

To further assess the generality of this strategy, we evaluated additional iLNP formulations. First, we tested a standard iLNP formulation based on the ionizable lipid D-Lin-MC3-DMA (MC3), which is used in approved siRNA therapeutics,^72, 73^ instead of the ALC-0315-based iLNPs used above. MC3-based iLNPs exhibited a hydrodynamic diameter of 105.4 ± 6.5 nm, a PDI of 0.09 ± 0.01, a ζ potential of 0.3 ± 0.8 mV, and an encapsulation efficiency of 82.0 ± 3.1 % (mean ± standard deviation; **Table S1**). Consistent with results obtained using ALC-0315-based iLNPs, 2-arm-PEG-oligocation pre-injection significantly reduced hepatic expression by 41-fold and increased splenic expression by 2.8-fold (**Figure S3A**), resulting in an improvement in spleen selectivity exceeding 100-fold (**Figure S3B**).

Up to this point, we used 2-arm-PEG-OligoLys, for which extensive characterization data are available from previous studies.^62, 63^ We next evaluated the feasibility of substituting this material with 2-armed PEG-oligoornithine (2-arm-PEG-OligoOrn), composed of 40 kDa PEG chains in each arm and approximately 20 ornithine units. Similar functionality is anticipated, as OligoOrn differs from OligoLys only in possessing one less methylene group in the side chain spacer (**Scheme 2**). To confirm this, we first examined the intrahepatic behavior of 2-arm-PEG-OligoOrn. Safe and effective LSE wall stealth coating requires efficient attachment of 2-arm-PEG-oligocations to the LSE wall followed by gradual detachment to avoid prolonged disruption of liver function. In previous work, 2-arm-PEG-OligoLys fulfilled these criteria by persisting in the LSE wall within several hours and subsequently undergoing biliary clearance.^62^ Similarly, IVRT-CLSM imaging revealed that 2-arm-PEG-OligoOrn efficiently and transiently attached to the LSE wall, with biliary excretion initiating approximately 3 h after injection (**Figure S4**), satisfying both safety and efficacy requirements. Furthermore, when combined with ALC-0315-based iLNPs, pre-injection of 2-arm-PEG-OligoOrn reduced hepatic expression by 28-fold, increased splenic expression by 2.6-fold (**Figure S5A**), and enhanced spleen selectivity by more than 100-fold (**Figure S5B**). Collectively, these results highlight the utility of 2-arm-PEG-OligoOrn for LSE wall stealth coating, yielding results similar to those observed with 2-arm-PEG-OligoLys.

### LSE wall stealth coating enhances vaccination efficacy after systemic iLNP injection

The data obtained thus far demonstrate the feasibility of both 2-arm-PEG-OligoLys and 2-arm-PEG-OligoOrn for mitigating hepatic accumulation of iLNPs. Among these candidates, we selected 2-arm-PEG-OligoOrn for subsequent applications in vaccination and immunotherapy, as it has already advanced toward clinical translation as a delivery carrier for siRNA or ASOs.^64–66^

Stealth coating of the LSE wall significantly enhanced protein expression from systemically administered iLNPs in the spleen (**Figures 4A, B, S1-3, S5**), particularly in splenic DCs (**Figure 4C, G**). This improvement suggests strong potential for mRNA vaccine applications, prompting us to evaluate the impact of this strategy on vaccination efficacy. Systemic administration of mRNA vaccines is predominantly employed in cancer vaccine strategies, aimed at inducing robust cellular immune responses.^17–22^ Accordingly, we focused on assessing cellular immunity, beginning with an enzyme-linked immunospot (ELISpot) assay. Mice were vaccinated twice with mRNA encoding *ovalbumin* (*OVA*) as a model antigen, with a 2-week interval between doses. ELISpot analysis was performed 1 week after the booster vaccination (**Figure 5A**). For each vaccination, ALC-0315-based iLNPs loaded with *OVA* mRNA were administered intravenously 5 min after systemic injection of 2-arm-PEG-OligoOrn. Notably, stealth coating significantly increased the number of antigen-reactive interferon γ (IFN- γ)-positive spots by approximately 2.5-fold, indicating enhanced cellular immune responses (**Figure 5B**).

**Figure 5.**
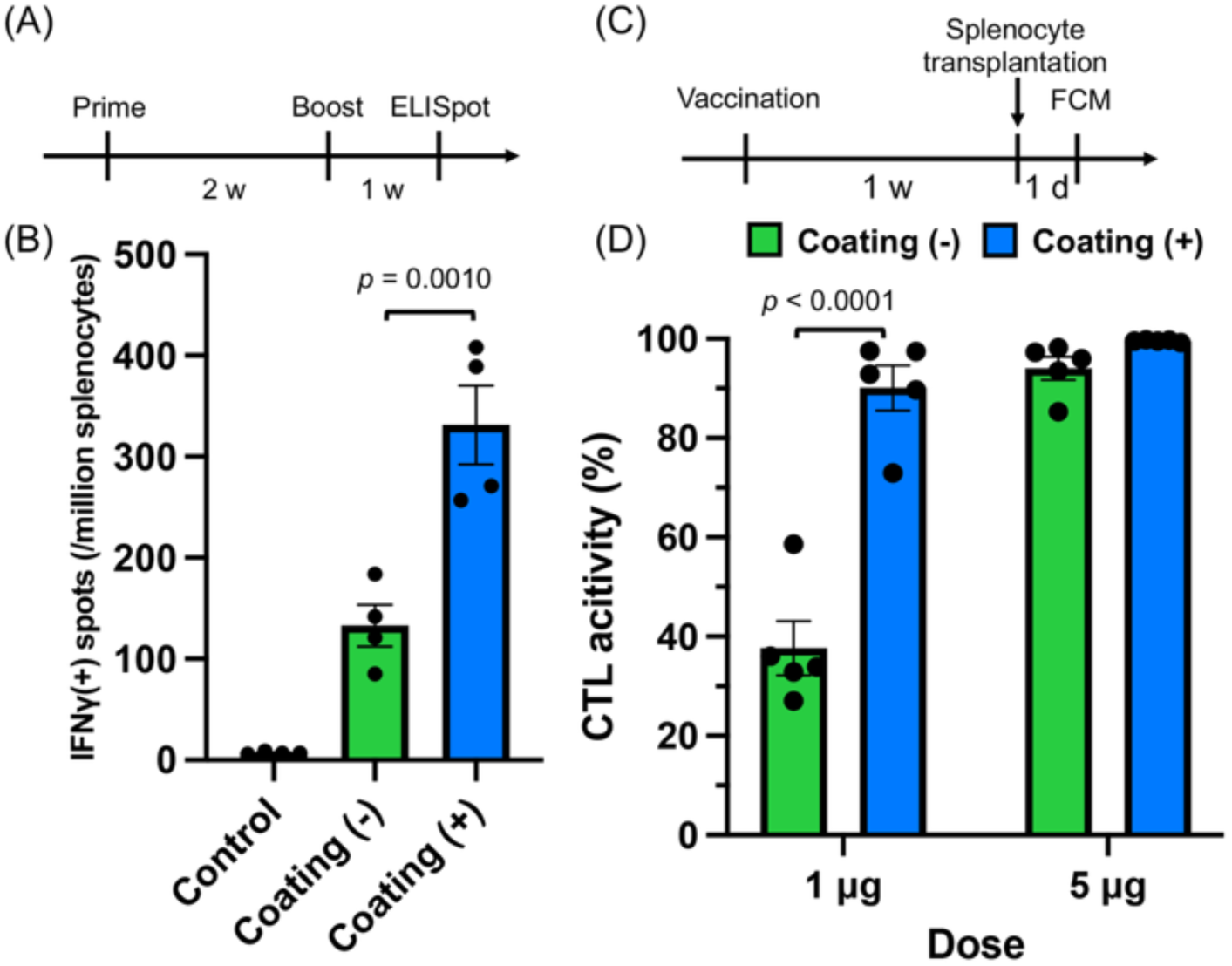
LSE wall stealth coating in systemic iLNP vaccination. ALC-0315-based iLNPs encapsulating *ovalbumin* (*OVA*) mRNA were intravenously administered with or without pre-injection of 2-arm-PEG-OligoOrn 5 min beforehand. (A, B) Mice were vaccinated twice at a 2-week interval, and ELISpot assays were performed 1 week after the booster. *n* = 4. (C, D) For the *in vivo* CTL assay, mice were vaccinated once. Splenocytes pulsed with an epitope peptide were transferred 7 days later, and target-cell elimination was analyzed by flow cytometry 1 day after transfer. *n* = 5. Data are mean ± SEM. Statistical analyses used one-way ANOVA with Tukey’s post hoc test (B) and two-way ANOVA with Šídák’s post hoc test (D).

To further evaluate functional cellular immunity, we conducted an *in vivo* CTL assay to assess the capacity of CTLs to eliminate antigen-expressing target cells, a critical function for effective cancer vaccination.^74^ In this assay, target splenocytes expressing the antigen were transferred into vaccinated mice, and target-cell elimination efficiency was quantified. Mice received systemic injections of 2-arm-PEG-OligoOrn followed by *OVA* mRNA-loaded iLNPs with a 5-min interval, and CTL activity was assessed one week later (**Figure 5C**). Mice treated with LSE wall stealth coating exhibited significantly enhanced CTL activity compared with those without coating at an mRNA dose of 1 μg (**Figure 5D**). These findings demonstrate that LSE wall stealth coating improves the efficacy of systemically administered iLNP vaccines, likely by enhancing protein expression efficiency in the spleen (**Figure 4**). Additionally, improved vaccine efficacy might also be attributed to reduced antigen expression in liver cells (**Figure 4**), which could limit activation of hepatic immune tolerance mechanisms.^49–51^

### LSE wall stealth coating reduces liver accumulation of locally injected iLNPs while enhancing cellular immunity induction

We next investigated the utility of LSE wall stealth coating in the context of locally administered mRNA iLNP vaccines, which present safety concerns due to unintended antigen expression in the liver. Following intramuscular injection, iLNPs can leak into systemic circulation and accumulate in the liver,^33, 35–37^ where ectopic antigen expression may trigger autoimmune hepatitis.^46, 47^ Indeed, a prior report described autoimmune hepatitis following COVID-19 mRNA iLNP vaccination and demonstrated infiltration of CD8^+^ T cells targeting the spike protein in the liver.^48^

To evaluate the effects of LSE wall stealth coating on locally delivered iLNPs, mice received systemic pre-injection of 2-arm-PEG-OligoOrn 5 min before intramuscular administration of ALC-0315-based iLNPs loaded with *luciferase* mRNA. Remarkably, stealth coating reduced luciferase expression in the liver by nearly 10-fold (**Figure 6A**). Importantly, protein expression was preserved at the injection site in skeletal muscle, as well as in draining lymph nodes and the spleen (**Figure 6A**), all of which are critical sites for effective mRNA vaccination.^12, 16, 25, 75^ In contrast to systemic iLNP delivery, LSE wall coating did not enhance protein expression efficiency in the spleen following intramuscular administration. This difference might reflect reduced systemic exposure to iLNPs after local injection, as a substantial fraction of iLNPs likely remains within the muscle tissue or accumulates in draining lymph nodes, as suggested by strong luciferase expression in these tissues (**Figure 6A**). Under these conditions, fewer iLNPs are redirected from the liver to the spleen, which may explain the limited impact of LSE wall stealth coating on splenic mRNA delivery after intramuscular injection.

**Figure 6.**
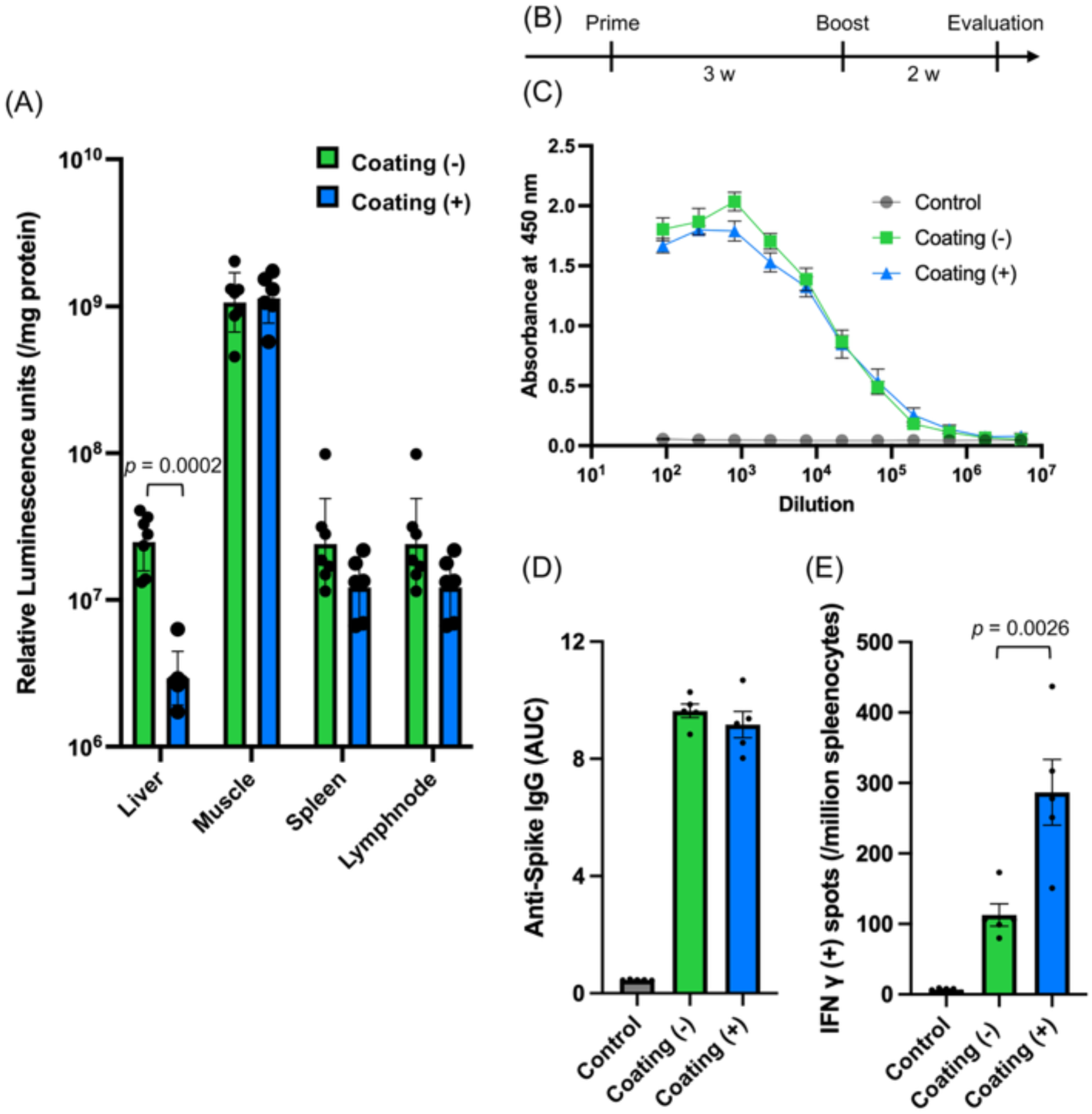
LSE wall stealth coating in intramuscular iLNP vaccination. ALC-0315-based iLNPs encapsulating *luciferase* mRNA or SARS-CoV-2 *spike* mRNA were injected intramuscularly with or without pre-injection of 2-arm-PEG-OligoOrn 5 min beforehand. (A) Luciferase expression in each organ 4 h after iLNP injection. *n* = 7 for coating (-), *n* = 6 for coating (+). Statistical analysis used an unpaired two-tailed Student’s *t*-test. (B) Vaccination schedule (two doses at a 3-week interval). (C, D) Spike-specific antibody measurement and (E) ELISpot analysis conducted 2 weeks after the booster. (C) Antibody titration curves. (D) Area under the curve derived from (C). *n* = 5. Data are mean ± SEM. Statistical analyses used one-way ANOVA followed by Tukey’s post hoc test.

We subsequently assessed the effect of LSE wall stealth coating on the efficacy of locally administered mRNA vaccines encoding the SARS-CoV-2 spike protein. Mice received two vaccinations at a 3-week interval, each consisting of systemic pre-injection of 2-arm-PEG-OligoOrn followed 5 min later by intramuscular iLNP administration (**Figure 6B**). Two weeks after the booster, enzyme-linked immunosorbent assay (ELISA) revealed comparable anti-spike IgG levels in mice with and without 2-arm-PEG-OligoOrn pre-injection (**Figures 6C, D**). Notably, however, ELISpot analysis demonstrated significantly enhanced cellular immune responses in mice receiving LSE wall stealth coating (**Figure 6E**). Unlike systemically administered vaccines (**Figure 5**), the enhanced cellular immunity observed after local vaccination cannot be explained by increased antigen expression in the spleen, which was similar with and without 2-arm-PEG-OligoOrn pre-injection. Instead, reduced antigen expression in the liver could contribute to improved vaccine efficacy by preventing activation of immune tolerance pathways, as the liver plays a central role in immune tolerance.^49–51^ From a safety perspective, this strategy effectively minimizes hepatic accumulation of locally administered iLNP vaccines without compromising vaccination efficacy. LSE wall stealth coating could therefore be particularly beneficial for individuals with pre-existing liver disease or autoimmune conditions. Notably, pro-inflammatory responses to iLNPs in the liver are amplified under pre-existing inflammatory conditions.^40^ These factors should be carefully considered when selecting patient populations in future translational studies to maximize the benefit of LSE wall stealth coating.

### LSE wall stealth coating mitigates safety concerns of intratumoral cytokine mRNA therapy

Beyond vaccines, cancer immunotherapy involving intratumoral administration of cytokine mRNA^41–45^ also raises safety concerns related to off-target protein expression in the liver. Systemic leakage of iLNPs from tumors and their subsequent accumulation in the liver can trigger adverse effects due to ectopic cytokine expression.^34, 39^ In this study, we applied 2-arm-PEG-OligoOrn pre-injection to address this issue, using intratumoral delivery of *interleukin 12* (*IL-12*) mRNA to melanoma as a model system.

Specifically, ALC-0315-based iLNPs loaded with *IL-12* mRNA were injected directly into established B16F10 melanoma, with or without systemic pre-injection of 2-arm-PEG-OligoOrn 5 min beforehand. Tumor growth was significantly suppressed (**Figure 7A**), and mouse survival was prolonged (**Figure 7B**) irrespective of 2-arm-PEG-OligoOrn pre-injection. To assess safety at this therapeutically effective iLNP dose, IL-12 protein levels were quantified in the tumor, liver, spleen, and serum 24 hours after intratumoral iLNP administration (**Figures 7C-F**). Pre-injection of 2-arm-PEG-OligoOrn did not affect IL-12 expression in the tumor (**Figure 7C**). Notably, intratumoral *IL-12* mRNA administration without 2-arm-PEG-OligoOrn pre-injection significantly increased IL-12 protein levels in the liver (**Figure 7D**), likely because of leakage of iLNPs from the tumor into systemic circulation and subsequent hepatic accumulation. In contrast, 2-arm-PEG-OligoOrn pre-injection reduced hepatic IL-12 concentrations to levels comparable to untreated controls, indicating effective prevention of liver accumulation of leaked iLNPs. Consistent with this observation, 2-arm-PEG-OligoOrn pre-injection also significantly reduced systemic IL-12 exposure, as reflected by decreased serum IL-12 levels (**Figure 7F**). Even in the absence of 2-arm-PEG-OligoOrn pre-injection, intratumoral *IL-12* mRNA administration did not elevate IL-12 levels in the spleen (**Figure 7E**), possibly because only a limited fraction of iLNPs entered systemic circulation following intratumoral administration. Collectively, these findings demonstrate that LSE wall stealth coating effectively mitigates systemic inflammatory risks associated with intratumoral cytokine mRNA therapy using iLNPs while maintaining antitumor efficacy.

**Figure 7.**
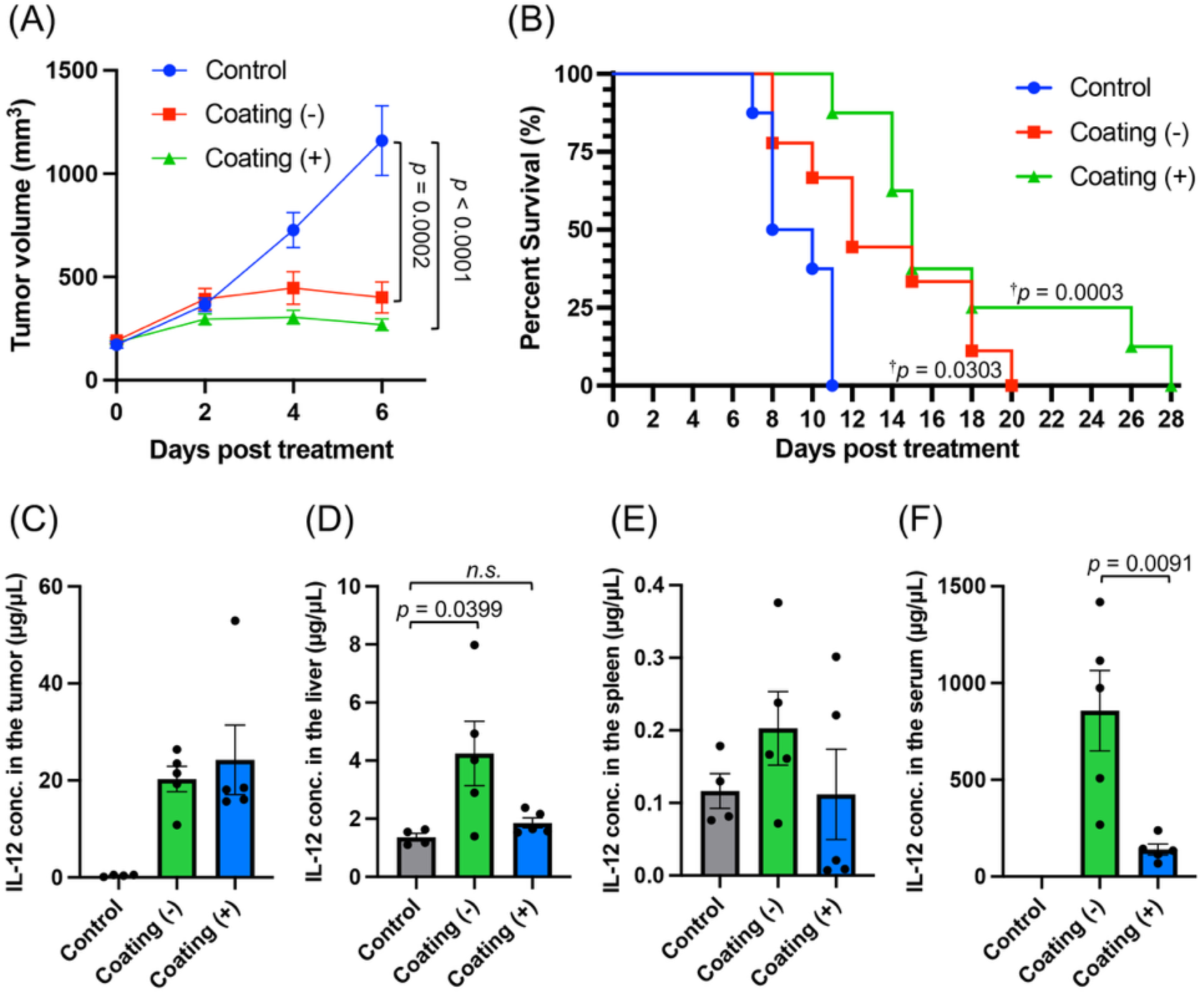
LSE wall stealth coating in intratumoral cytokine mRNA therapy. Mice were subcutaneously inoculated with B16F10 melanoma cells. Once tumors reached a defined size, ALC-0315-based iLNPs encapsulating *IL-12* mRNA were injected intratumorally with or without pre-injection of 2-arm-PEG-OligoLys 5 min beforehand. (A) Tumor growth curves. (B) Survival analysis. *n* = 8. (C-F) Safety assessment. IL-12 protein levels in the tumor (C), liver (D), spleen (E), and serum (F) were quantified by ELISA 24 h post-iLNP injection. *n* = 5. Data are mean ± SEM. Statistical analyses used one-way ANOVA followed by Tukey’s post hoc test in (A, D, F) and a log-rank test in (B; †: vs. Control).

## CONCLUSION

This study demonstrates that LSE wall stealth coating using 2-arm-PEG-oligocations effectively reduces hepatic accumulation of mRNA-loaded iLNPs and the resulting off-target protein expression in the liver. Importantly, LSE wall coating does not compromise protein expression efficiency in the spleen or lymph nodes, which is critical for effective mRNA vaccination. This strategy addresses two major challenges associated with hepatic iLNP accumulation. First, when extrahepatic tissues are the intended targets, liver sequestration of iLNPs limits their delivery to these sites. In this study, LSE wall coating redirected systemically administered iLNPs from the liver to the spleen, thereby enhancing vaccine efficacy. This approach may be broadly applicable to diverse iLNP systems designed to target extrahepatic cell types and organs, including T cells, hematopoietic stem cells, tumors, and the brain, which are currently the focus of intensive research efforts.^76–81^ Second, LSE wall stealth coating mitigates adverse effects arising from unintended hepatic protein expression following iLNP administration. As demonstrated here, this strategy reduced hepatic cytokine expression after intratumoral administration of cytokine mRNA and minimized antigen expression in the liver following systemic or intramuscular administration of iLNP vaccines. In doing so, it improves the safety profile of existing mRNA therapeutics and vaccines.

Importantly, all vaccine and immunotherapy applications in this study employed clinically relevant materials, including 2-arm-PEG-OligoOrn currently under clinical evaluation^64–66^ and clinically approved ALC-0315-based iLNP formulations. For clinical translation, 2-arm-PEG-oligocations have already been produced under Good Manufacturing Practice (GMP) conditions and have passed rigorous preclinical safety assessments. Furthermore, they recently demonstrated safety in a phase I trial up to the initially determined maximum tolerated dose.^82^ This high degree of clinical translatability may offer a substantial advantage over alternative approaches aimed at reducing hepatic iLNP accumulation, such as developing entirely new iLNP designs.^39, 83, 84^ At the same time, combining LSE wall stealth coating with existing strategies, such as the use of novel iLNP formulations or incorporation of hepatocyte-specific microRNA target sequences within untranslated regions,^34, 42^ represents a promising direction. Because the LSE wall coating strategy operates through a distinct mechanism, it has strong potential to synergize with these complementary approaches, further enhancing the safety and efficacy of mRNA-based vaccines and immunotherapies.

## EXPERIMENTAL SECTION

### Synthesis of 2-arm-PEG-OigoLys and 2-arm-PEG-OligoOrn

2-arm-PEG-OligoLys was synthesized as reported previously.^62, 63^ Briefly, 2-arm-PEG-NH_2_ (0.37 g, 5 μmol) and *N*^ε^-trifluoroacetyl-L-lysine-*N*-carboxyanhydride (L-Lys(TFA)-NCA, 35 mg, 130 μmol) were reacted in DMF (3.6 mL for 2-arm-PEG-NH_2_ and 2.5 mL for L-Lys (TFA)-NCA, respectively) containing 1 M thiourea under argon at 25 °C for 48 h. The reaction mixture was reprecipitated twice in diethyl ether and vacuum dried. TFA groups of the resulting 2-arm-PEG-OligoLys(TFA) (200 mg in 20 mL of methanol) were removed by treatment with 1 M NaOH(aq) (2 mL) at 35 °C for 16 h. The solution was purified by dialysis (MWCO 12 kDa) against 0.01 M HCl, 0.001 M HCl, and deionized water, followed by filtration (0.22 µm) and lyophilization to yield the HCl salt of 2-arm-PEG-OligoLys. 2-arm-PEG-OligoOrn was synthesized in an analogous manner using L-Orn(TFA)-NCA, with deprotection performed at 25 °C and dialysis at 4 °C.^85^ The degrees of polymerization (DP) of OligoLys and OligoOrn were determined by ^1^H NMR (D_2_O, 25 °C; JEOL JNM-ECS400) and found to be 20 for OligoLys and 19 for OligoOrn, respectively. Calculations were based on integration ratios of PEG methylene protons (δ = 3.5–4.0 ppm) to side-chain methylene protons of Lys (δ = 1.2–2.0 ppm) and Orn (δ = 1.6–2.0 ppm). Molecular weight dispersity (*Đ*) was determined by aqueous size-exclusion chromatography (SEC) calibrated with PEG standards using a JASCO HPLC LC-2000 system equipped with a Superdex 200 Increase 10/300 GL column (GE Healthcare, Chicago, IL) and ultraviolet (UV) (220 nm) and refractive index (RI) detectors. Elution was performed with 10 mM acetic acid (pH 3.3) containing 500 mM NaCl at a flow rate of 0.5 mL/min at 25 °C. Both polymers exhibited unimodal elution profiles with *Đ* = 1.04.

### Fluorescence labeling of 2-arm-PEG-OigoLys and 2-arm-PEG-OligoOrn

2-arm-PEG-OligoLys(TFA) (20 mg in 4 mL of 0.1 M sodium bicarbonate buffer, pH 8.5) was reacted with Alexa Fluor^TM^ 647 (A647) NHS ester (44 μL of a 20 mg/mL DMF solution) for 48 h at 25 °C. The reaction mixture was dialyzed against deionized water and lyophilized, followed by TFA deprotection, as described above. The product was purified using a PD-10 column (1 M NaCl) and dialyzed against deionized water to yield A647–2-arm-PEG-OligoLys. Other labeled polymers (A594-, A555-, and A450–2-arm-PEG-OligoLys) and A647–2-arm-PEG-OligoOrn were synthesized using the corresponding NHS ester dyes following the same procedure.

### mRNA and iLNP preparation

*Luciferase* mRNA prepared in-house as described previously^86^ was used for **Figures 4A, B, S1, and S2**. *Luciferase* mRNA for **Figures S3 and S5**, *ovalbumin* (*OVA*) mRNA, and *Cre recombinase* mRNA without nucleoside modification, and CleanCap N1-methylpseudouridine-modified SARS-CoV-2 spike mRNA (identical in sequence to that in BNT162b2)^7^ were purchased from TriLink Biotechnologies (San Diego, CA, USA). *NanoLuc luciferase* mRNA used for **Figures 1B**, **6A** and single-chain *interleukin* (*IL*)*-12* mRNA^42^ with N1-methylpseudouridine modification were obtained from GenScript (Piscataway, NJ, USA). iLNPs were prepared by ethanol dilution methods from an aqueous sodium citrate buffer containing mRNA and an ethanol solution of four lipid components, as described previously for ALC-0315-based iLNPs^36^ and MC3-based iLNPs.^87^ The lipid molar ratios were ALC-0315: DSPC: cholesterol: PEG-lipid: ALC-0159 = 46.3: 9.4: 42.7: 1.6 for ALC-0315-based iLNPs and D-Lin-MC3-DMA: DSPC: cholesterol: PEG2000-DMG = 50:10:38.5:1.5 for MC3-based iLNPs. [Nitrogen in ionizable lipids (N)] / [phosphate in mRNA (P)] ratio was 6 for ALC-0315-based iLNPs and 5 for MC-3-based iLNPs. Dynamic light scattering (DLS) measurements were performed using a Zetasizer Nano-ZS ZEN3550 (Malvern Instruments, Worcestershire, UK) equipped with a 523 nm diode laser and a scattering angle of 173°. ζ-potential was measured using laser Doppler electrophoretic light scattering with the same instrument.

### Evaluation of polymer binding to iLNP

The interaction between A594–2-arm-PEG-OligoLys and iLNPs was analyzed by fluorescence correlation spectroscopy (FCS). Diffusion times were derived from the autocorrelation functions of the polymer in both its free state and in the presence of iLNPs. Measurements were conducted at 37 °C in 10 mM D-PBS, with a polymer concentration of 0.625 mg/mL and iLNPs loaded with 2.5 µg/mL mRNA. Data were acquired using a Zeiss LSM880 confocal microscope equipped with a C-Apochromat 40× water-immersion objective.

### Animal experiments

BALB/c and C57BL/6J mice were purchased from Charles River Laboratories Japan (Yokohama, Japan). All animal experiments were approved by the animal committees in Kyoto Prefectural University of Medicine (Kyoto, Japan), Institute of Science Tokyo (Tokyo, Japan), or Innovation Center of NanoMedicine, Kawasaki Institute of Industrial Promotion (Kawasaki, Japan).

### Biodistribution of 2-arm-PEG-oligocations

Thirty minutes after intravenous injection of Alexa594-labeled 2-arm-PEG-OligoLys (1.25 mg) via the tail vein, BALB/c mice (7w old, female) were euthanized. The liver, spleen, lung, kidney, heart, and brain were excised, briefly rinsed with PBS, blotted dry, weighed, and homogenized in Passive Lysis Buffer (Promega) using a Multi-Beads Shocker (Yasui-Kikai, Osaka, Japan). Homogenates were centrifuged to remove debris, and fluorescence intensity of the supernatants was measured using a Spark microplate reader (TECAN, Männedorf, Switzerland) with excitation at 590 nm and emission at 617 nm.

The percentage of injected dose per gram of tissue (% dose/g) was calculated by dividing the fluorescence intensity of each organ homogenate by tissue weight (g), normalizing to total administered fluorescence, and multiplying by 100. Total administered fluorescence was estimated by measuring the fluorescence intensity from a mixture of 1.25 mg of Alexa594-labeled 2-arm-PEG-OligoLys (corresponding to the injected dose) and organ homogenates from buffer-injected control mice.

### Biodistribution of mRNA-loaded iLNPs

Four hours after intravenous injection into BALB/c mice (7w old, female), the liver, spleen, lung, kidney, heart, and brain were harvested and homogenized in Buffer RLT (RNeasy Mini Kit, Qiagen, Hilden, Germany) containing 1% 2-mercaptoethanol using a Micro Smash MS-100 homogenizer (Tomy Digital Biology, Tokyo, Japan). Total RNA was extracted using the RNeasy Mini Kit, followed by enzymatic digestion of genomic DNA and complementary DNA (cDNA) synthesis by reverse transcription using the ReverTra Ace™ qPCR RT Master Mix with gDNA Remover (TOYOBO, Osaka, Japan). Quantitative PCR (qPCR) was performed using SYBR Green qPCR Master Mix on a QuantStudio™ 1 Real-Time PCR System (Thermo Scientific, Waltham, MA, USA). Relative mRNA levels were calculated using the 2^−ΔCt^ method, with β-actin as the endogenous reference gene. Primer sequences were GGCGTAATGATAGGCTCGCT and CAGATGCTCAAGGCCCTTCA for *NanoLuc luciferase* and CATTGCTGACAGGATGCAGAAGG and TGCTGGAAGGTGGACAGTGAGG for *β-actin*.

### IVRT-CLSM observation

BALB/c mice (7w old, female) were anesthetized with 2.5% isoflurane, and the liver was surgically exposed and partially immobilized under a coverslip. Intravital liver imaging was performed using a confocal laser-scanning microscope (A1R, Nikon Corp., Tokyo, Japan) mounted on an upright microscope (ECLIPSE FN1, Nikon Corp.). A 40× water-immersion objective was used, and the pinhole diameter was adjusted to obtain optical sections of 5 μm. Mice were intravenously injected with 1.25 mg of Alexa555–labeled 2-arm-PEG-OligoLys and 5 μg of luciferase mRNA–loaded, DiD-labeled ALC-0315–based iLNPs sequentially at a 5-min interval. Liver autofluorescence was excited at 488 nm and detected using a 525/50 nm bandpass filter. Alexa555 was excited at 561 nm and detected using a 595/50 nm bandpass filter, while DiD was excited at 640 nm and detected with a 700/75 nm bandpass filter.

For spleen imaging, intravital real-time multiphoton microscopy was employed. BALB/c mice (7w old, female) were anesthetized with 2.5% isoflurane, and the spleen was surgically exposed and partially immobilized under a coverslip. Imaging was performed using a multiphoton microscope (A1R MP, Nikon Corp.) equipped with a pulsed laser system (InSight DeepSee, Spectra-Physics, Santa Clara, CA, USA). A 20× water-immersion objective was used. Mice were intravenously injected with 1.25 mg of Alexa450–labeled 2-arm-PEG-OligoLys and 5 μg of *luciferase* mRNA–loaded DiD-labeled ALC-0315–based iLNPs sequentially at a 5-min interval. To identify iLNP entry sites in the spleen, 10 μg of Alexa Fluor488^TM^–labeled anti-F4/80 antibody (Invitrogen, Carlsbad, CA, USA) was injected via the tail vein to visualize red pulp macrophages. In this experiment, mice were sequentially injected with unlabeled 2-arm-PEG-OligoLys, followed by 5 μg of *luciferase* mRNA–loaded DiD-labeled iLNPs at a 5-min interval. Excitation was performed at 820 nm, and emitted fluorescence signals were collected using bandpass filters of 450/70 nm for Alexa450, 525/50 nm for Alexa488, and 700/75 nm for DiD. Image acquisition and processing were performed using NIS-Elements software (Nikon Corp.).

### Luciferase expression assays

BALB/c mice (7w old, female) were intravenously injected with 1.25 mg of 2-arm-PEG-OligoLys or 2-arm-PEG-OligoOrn, followed 5 min later by intravenous injection of 5 μg of *luciferase* mRNA-loaded iLNPs. Organs were harvested 4 h or 24 h post-injection and homogenized in Passive Lysis Buffer (Promega, Madison, WI, USA) using a Micro Smash MS-100 homogenizer (Tomy Digital Biology, Tokyo, Japan). Luminescence was measured using a Lumat3 LB9508 luminometer (Berthold Technologies, Bad Wildbad, Germany) and normalized to protein amount determined with the Micro BCA Protein Assay Kit (Thermo Fisher Scientific, Waltham, MA, USA).

### tdTomato expression assays using transgenic mice

B6.Cg-Gt(ROSA)26Sortm9(CAG-tdTomato)Hze/J (Ai9) mice (Jackson Laboratory, Bar Harbor, ME, USA) were sequentially injected via the tail vein with 1.25 mg of 2-arm-PEG-OligoLys and 5 μg of *Cre* mRNA loaded in ALC-0315-based iLNPs at a 5-min interval. Seventy-two hours post-injection, the liver was perfused sequentially with buffer and collagenase solutions to dissociate cells. The composition of these solutions and detailed procedures were described previously.^88^ Collected cells were centrifuged at 50 × g for 4 min at 4 °C. The pellet was used for hepatocyte isolation, while the supernatant was used for isolation of LSECs and KCs. Hepatocytes were isolated by repeated low-speed centrifugation (50 × g, 4 min, 4 °C, twice). LSECs and KCs were isolated from the supernatant using CD146 MicroBeads for LSECs and Anti-F4/80 MicroBeads UltraPure for KCs, respectively (MACS cell separation, Miltenyi, Bergisch Gladbach, Germany). For DC analysis, splenocytes were stained with Alexa Fluor^TM^ 488-conjugated anti-mouse CD11c antibody (BioLegend, San Diego, CA, USA). Flow cytometric analysis was performed using an LSRFortessa X-20 cell analyzer (BD Biosciences, Franklin Lakes, NJ, USA).

### Vaccination by systemic iLNP injection

C57BL/6J mice (7w old, female) were intravenously injected with 1.25 mg of 2-arm-PEG-OligoOrn, followed 5 min later by 5 μg of *OVA* mRNA loaded in ALC-0315-based iLNPs. For ELISpot analysis, mice were vaccinated twice at a two-week interval, and splenocytes were collected 1 w after the booster. Splenocytes were seeded at a density of 2.5× 10^5^ cells/well in anti-IFNγ ELISpot 96 well plates and stimulated with 0.025 μg/well PepTivator^®^ Ovalbumin epitope mix (Nordrhein-Westfalen, Germany). After overnight incubation, plates were processed according to the manufacturer’s protocol and analyzed using an ELISpot reader (AID GmbH, Germany). *In vivo* CTL assays were performed as described previously.^36, 89^ Briefly, splenocytes from donor mice were divided into two populations: one was pulsed with SIINFEKL peptide (Toray, Tokyo, Japan) and labeled with 5 mM CFSE (Molecular Probes, Eugene, OR, USA), while the other was labeled with 0.5 mM CFSE without peptide pulsing. Equal numbers of both populations (1 × 10^7^ cells in total) were intravenously injected into mice 7 days-post vaccination. Twenty-four hours later, splenocytes were collected and analyzed by flow cytometry (BD LSRFortessa^TM^ X-20).

### Intramuscular injection of iLNPs

BALB/c mice (7w old, female) were intravenously injected with 1.25 mg of 2-arm-PEG-OligoOrn. Five minutes later, ALC-0315-based iLNPs containing 5 μg of *luciferase* or SARS-CoV-2 *spike* mRNA were injected into the femoral muscles. Luciferase expression was analyzed 4 h post-injection as described in the section “Luciferase expression assays”. For vaccination studies, mice were immunized twice at a three-week interval. Plasma and splenocytes were collected 2 w after the booster dose. ELISpot assays were performed as described in the section “Vaccination by systemic iLNP injection” except that splenocytes were stimulated overnight with 0.2 μg/well PepMix™ SARS-CoV-2 (S) (JPT Peptide Technologies, Berlin, Germany). For antibody measurements, 50 μL/well of recombinant spike protein (2 μg/mL in 50 mM carbonate buffer, pH=9.6) was coated onto clear flat-bottom immuno nonsterile 96-well plates (Thermo Fisher Scientific) by overnight incubation at 4 °C. Plates were washed 3 times with PBS containing 0.5% w/v Tween 20 (PBS-T) and blocked with diluent (1% BSA, 2.5 mM EDTA in PBS-T) at room temperature for 1 h. After removing the solution, plasma samples diluted in blocking buffer were added (50 μL/well) and incubated overnight at 4 °C. After washing 3 times with PBS-T, plates were incubated with goat anti-mouse HRP-conjugated secondary antibody (R&D Systems, Minneapolis, MN, USA; 1:8000 dilution) for 2 h at room temperature. Detection was performed using 3,3′,5,5′-tetramethylbenzidine (TMB) substrate (100 μL), and the reaction was stopped with 2 N sulfuric acid (50 µL). Absorbance was measured at 450 nm using a microplate reader (Infinite^®^ 200PRO F Nano+, Tecan, Männedorf, Switzerland).

### Intratumoral injection

B16F10 melanoma tumors were established by subcutaneous inoculation into the right flank of C57BL/6J mice (6-week-old females). Specifically, mice were injected with 1 × 10^6^ cells suspended in 100 µL of DMEM containing 1% Matrigel. Tumor size was measured using a digital caliper and expressed as tumor volume calculated according to the formula: Volume = 1/2 × (long axis) × (short axis)^2^

When tumors reached a volume of 90–100 mm³, mice received a single intratumoral injection of iLNPs encapsulating 5 μg of *IL-12* mRNA. Injections were performed using a 29-gauge needle inserted into the tumor core under isoflurane anesthesia, following an intravenous pre-injection of 1.25 mg of 2-arm-PEG-OligoOrn administered 5 min beforehand. Mice were euthanized when tumor volume exceeded 1,500–2,000 mm³, when tumor ulceration occurred, or when body weight loss exceeded 20%, in accordance with humane endpoint criteria.

IL-12 protein levels were analyzed 24 h after intratumoral administration of *IL-12* mRNA-loaded iLNPs. Tumor, liver, and spleen tissues were harvested and homogenized in lysis buffer containing protease inhibitors, followed by centrifugation at 12,000 × g for 10 min at 4 °C to remove debris. Serum samples were prepared by centrifugation at 10,000 × g for 10 min. IL-12 concentrations in tissue lysates and serum were quantified using a Quantikine™ Mouse IL-12 p70 ELISA kit in combination with the DuoSet™ Ancillary Reagent Kit 2 (R&D Systems). IL-12 levels were normalized to total protein content, which was determined using the Pierce™ BCA Protein Assay Kit (Thermo Fisher Scientific).

## AUTHOR INFORMATION

### Corresponding Authors

Kazunori Kataoka: Innovation Center of NanoMedicine (iCONM), Kawasaki Institute of Industrial Promotion, 3-25-14 Tonomachi, Kawasaki-ku, Kawasaki 210-0821, Japan

Satoshi Uchida: Department of Advanced Nanomedical Engineering, Medical Research Laboratory, Institute of Integrated Research, Institute of Science Tokyo, 1-5-45 Yushima, Bunkyo-ku, Tokyo 113-8510, Japan; Innovation Center of NanoMedicine (iCONM), Kawasaki Institute of Industrial Promotion, 3-25-14 Tonomachi, Kawasaki-ku, Kawasaki 210-0821, Japan; Medical Chemistry, Graduate School of Medical Science, Kyoto Prefectural University of Medicine, 1-5 Shimogamohangi-cho, Sakyo-ku, Kyoto 606-0823, Japan; Pandemic Preparedness, Infection and Advanced Research Center (UTOPIA), The University of Tokyo, 4-6-1, Shirokanedai, Minato-ku, Tokyo 108-0071, Japan;

### Author Contributions

The manuscript was written through contributions of all authors. All authors have given approval to the final version of the manuscript. ‡These authors contributed equally.

Conceptualization: S.U.; Methodology: K.T.; Investigation: A.D., B.C., L.B.T.N., K.T., M.M.M., X.L., T.A.T., S.F.; Writing - Original Draft: S.U.; Writing - Review & Editing: K.K., S.U.; Supervision: N.Q, J.I, J.N., Y.M., M.O., K.K., S.U.

### Notes

D.A, M.M.M., K.K., and S.U. have filed a patent application related to this study, and NANO MRNA Co., Ltd. holds a right to the patent. M.M.M. is an employee of NANO MRNA Co., Ltd.

## ACKNOWLEDGMENT

We would like to thank Sachi Ibuki (Kyoto Prefectural University of Medicine), Erika Mochizuki, Nao Horii, Katsunori Iwasa, Reiko Shiratori (Institute of Science Tokyo), Yuki Sato (iCONM), and Yumi Fujii (NANO MRNA Co., Ltd.) for their technical assistance.

