## Supplementary Information for "2-arm-PEG-oligocations transiently shield the liver sinusoids to mitigate off-target hepatic expression of mRNA lipid nanoparticles"

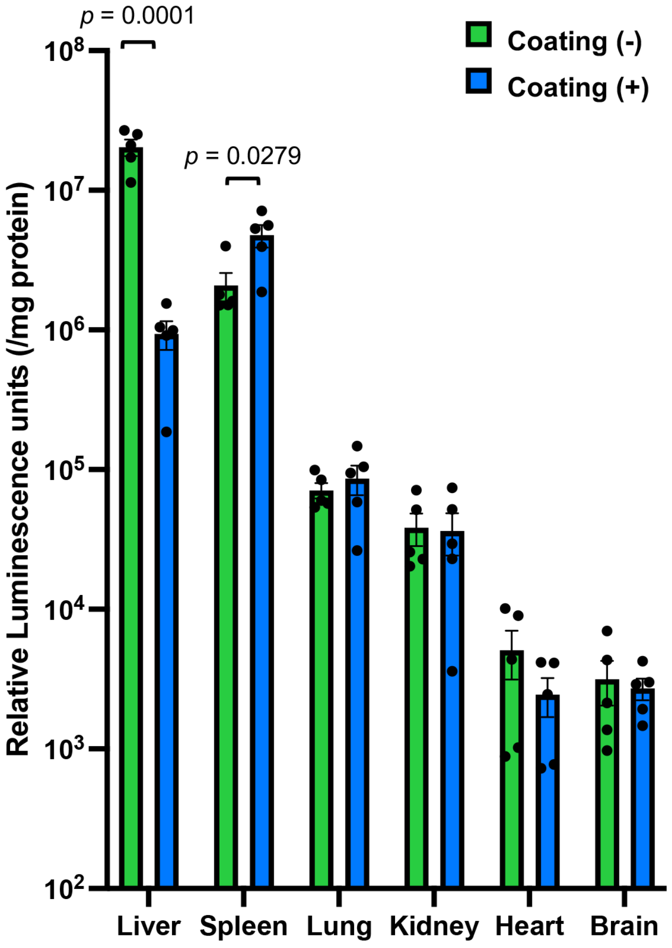


Supplementary Figure S1. Luciferase expression in organs following systemic injection of ALC-0315-based iLNPs with or without pre-injection of 2-arm-PEG-OligoLys 5 min beforehand. Data are reproduced from Figure 4A and displayed on a logarithmic scale. $n = 5$. Data are presented as mean ± SEM. Statistical analysis was performed using an unpaired two-tailed Student's $t$-test.

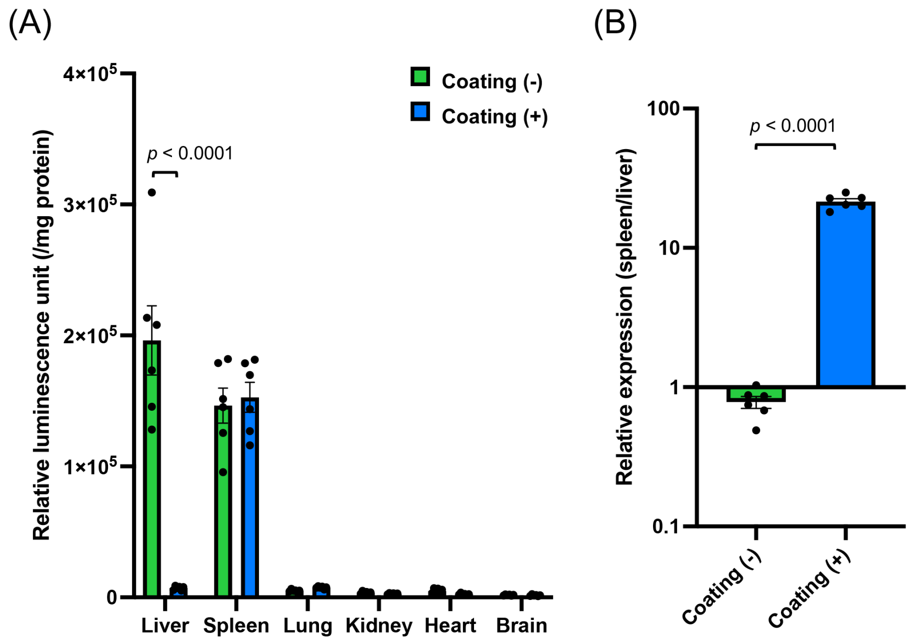


Supplementary Figure S2. Effects of LSE wall stealth coating on protein expression 24 hours post-injection. Luciferase expression was quantified 24 h after intravenous administration of ALC-0315-based iLNPs with or without pre-injection of 2-arm-PEG-OligoLys 5 min beforehand. (A) Luciferase expression in individual organs. (B) Relative spleen-to-liver expression normalized to organ weight. $n = 6$. Data are presented as mean ± SEM. Statistical analysis was performed using an unpaired two-tailed Student's $t$-test.

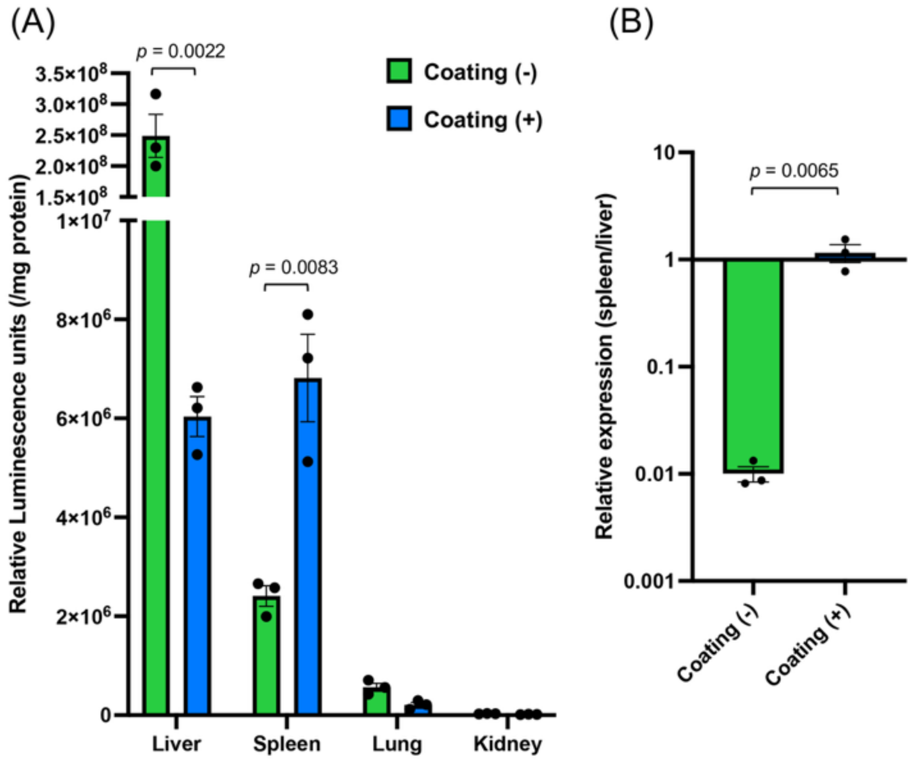


Supplementary Figure S3. Luciferase expression following systemic injection of MC-3-based iLNPs. MC-3-based iLNPs encapsulating *luciferase* mRNA were intravenously injected with or without pre-injection of 2-arm-PEG-OligoLys 5 min beforehand. (A) Luciferase expression in each organ. (B) Relative spleen-to-liver expression normalized to organ weight. $n = 3$. Data are presented as mean ± SEM. Statistical analysis was performed using an unpaired two-tailed Student's $t$-test.

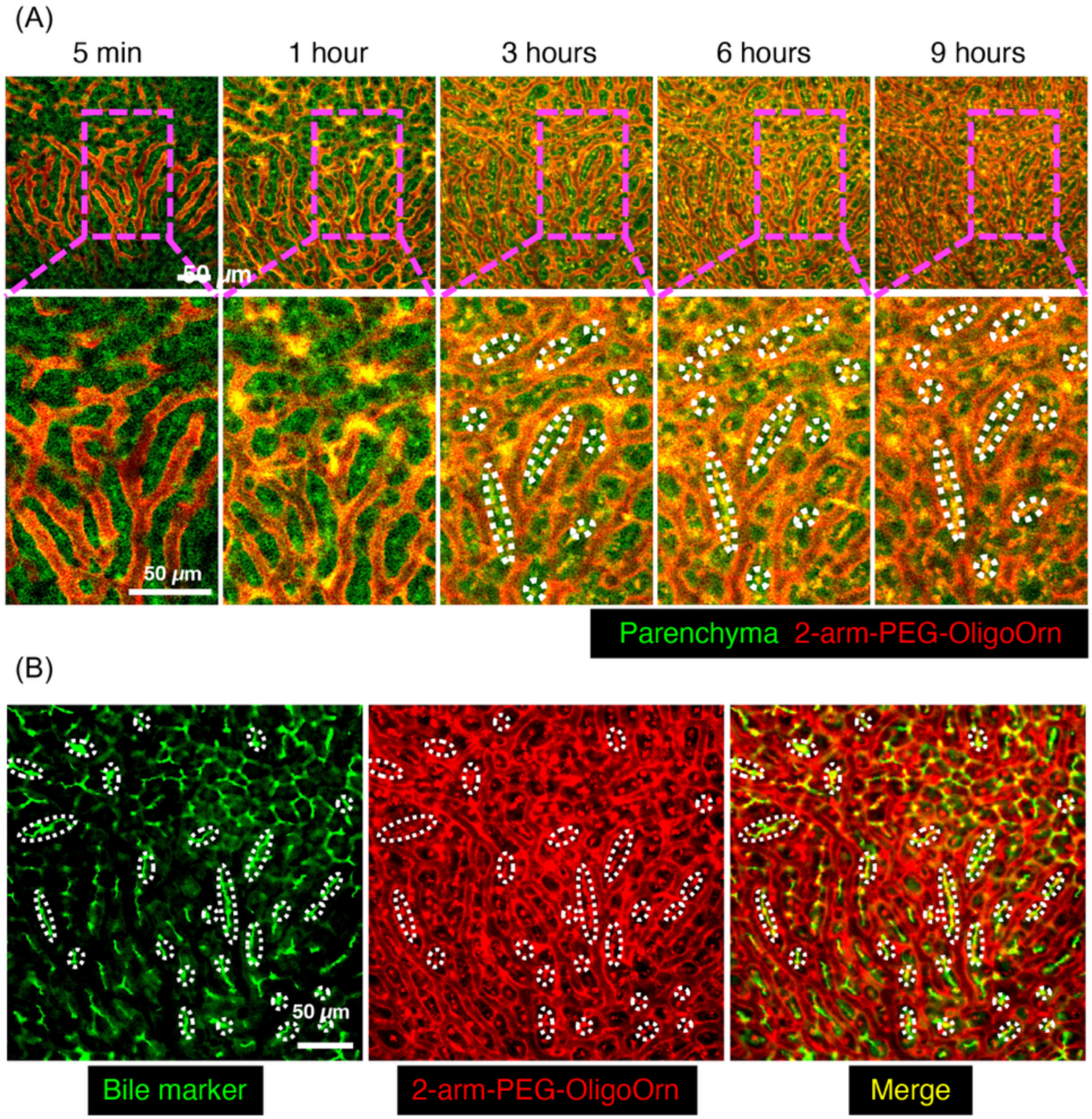


Supplementary Figure S4. Real-time observation of 2-arm-PEG-OligoOrn in the liver. Alexa647-labeled 2-arm-PEG-OligoOrn (red) was intravenously injected. (A) Liver parenchymal autofluorescence is shown in green. Images were acquired at the indicated time points. (B) Liver imaging 9 h post-injection, with bile canaliculi visualized in green using intravenous injection of 5-carboxyfluorescein. Bile canaliculi are outlined with white dotted lines.

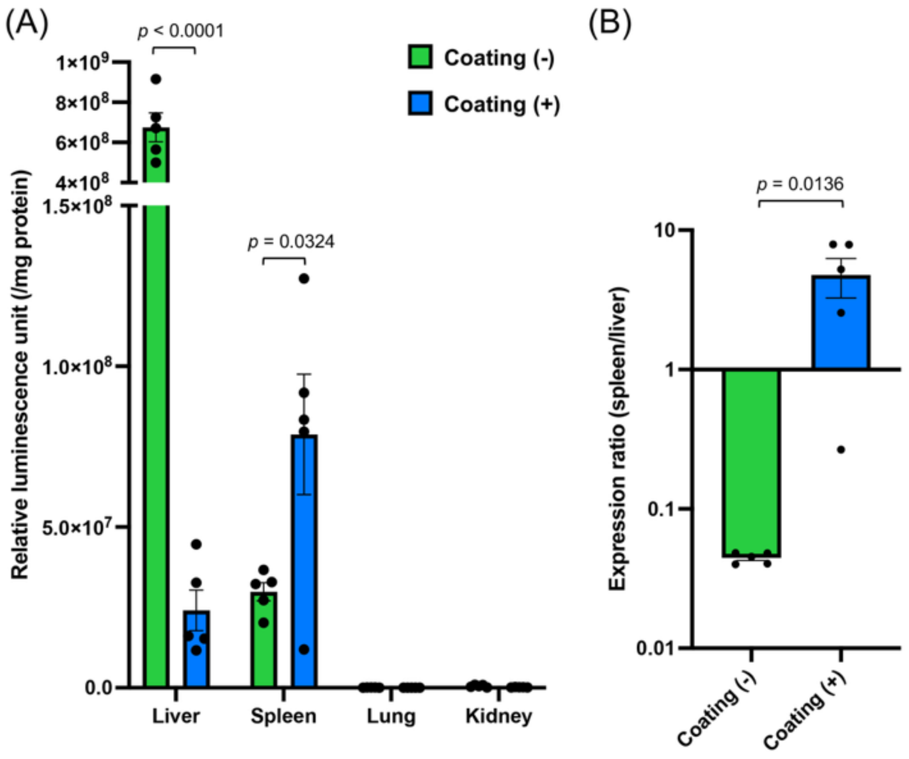


Supplementary Figure S5. Effect of pre-injection of 2-arm-PEG-OligoOrn on luciferase expression. ALC-0315-based iLNPs encapsulating *luciferase* mRNA were intravenously injected with or without pre-injection of 2-arm-PEG-OligoOrn 5 min beforehand. (A) Luciferase expression in individual organs. (B) Relative spleen-to-liver expression normalized to organ weight. $n = 5$. Data are presented as mean ± SEM. Statistical analysis was performed using an unpaired two-tailed Student's $t$-test.

Table S1. Characterization of iLNPs

| Ionizable lipids | Size (d., nm) | PDI | ζ-potential | Encapsulation efficiency (%) |
|---|---|---|---|---|
| ALC-0315 | 83.8 ± 0.7 | 0.09 ± 0.02 | -5.8 ± 1.7 | 95.7 ± 0.8 |
| MC-3 | 105.4 ± 6.5 | 0.09 ± 0.01 | 0.3 ± 0.8 | 82.0 ± 3.1 |

*Luciferase* mRNA-encapsulating iLNPs were analyzed. $n = 3$.

Table S2. Blood chemistry analysis 24 h after iLNP injection

| | AST (IU/L) | ALT (IU/L) | LDH (IU/L) | BUN (mg/dL) | CRE (mg/dL) |
|---|---|---|---|---|---|
| Control | 106 ± 10 | 97 ± 20 | 567 ± 113 | 31 ± 2 | 0.16 ± 0.02 |
| Coating (-) | 81 ± 6 | 66 ± 8 | 405 ± 124 | 30 ± 2 | 0.15 ± 0.01 |
| Coating (+) | 90 ± 16 | 65 ± 11 | 278 ± 33 | 27 ± 2 | 0.13 ± 0.02 |

$n = 4$. Plasma samples were analyzed using a DRI-CHEM 7000i system (Fujifilm, Tokyo, Japan). Abbreviations: IU, international unit; AST, aspartate aminotransferase; ALT, alanine aminotransferase; LDH, lactate dehydrogenase; BUN, blood urea nitrogen; Cre, creatinine.
